# Coupling cryo-electron tomography with mixed-scale dense neural networks reveals re-organization of the invasion machinery of *Toxoplasma gondii* upon ionophore-stimulation

**DOI:** 10.1101/2022.01.12.476068

**Authors:** Li-av Segev-Zarko, Peter D. Dahlberg, Stella Y. Sun, Daniël M. Pelt, Chi Yong Kim, Elizabeth S. Egan, James A. Sethian, Wah Chiu, John C. Boothroyd

## Abstract

Host cell invasion by intracellular, eukaryotic parasites within the phylum Apicomplexa, is a remarkable and active process involving the coordinated action of apical organelles and other structures. To date, capturing how these structures interact during invasion has been difficult to observe in detail. Here, we used cryogenic electron tomography to image the apical complex of *Toxoplasma gondii* tachyzoites under conditions that mimic resting parasites and those primed to invade through stimulation with calcium ionophore. Through the application of Mixed Scale Dense networks for image-processing, we developed a highly efficient pipeline for annotation of tomograms, enabling us to identify and extract densities of relevant subcellular organelles and accurately analyze features in 3D. The results reveal a dramatic change in the shape of the anteriorly located apical vesicle upon its apparent fusion with a rhoptry, that occurs only in the stimulated parasites. We also present information indicating that this vesicle originates from the vesicles that parallel the intraconoidal microtubules and that the latter two structures are linked by a novel tether. We show that a rosette structure previously proposed to be involved in rhoptry secretion is associated with apical vesicles beyond just the most anterior one. This result, suggesting multiple vesicles are primed to enable rhoptry secretion, may shed light on the mechanisms *Toxoplasma* employs to enable repeated invasion attempts. Using the same approach, we examine *Plasmodium falciparum* merozoites and show that they too possess an apical vesicle just beneath a rosette, demonstrating evolutionary conservation of this overall subcellular organization.

**Significance Statement:** Parasites in the phylum Apicomplexa are responsible for some of the most important parasitic diseases of humans, such as malaria and toxoplasmosis. Invasion by these obligatory, intracellular parasites depends on protein injection into the host cell. Using cryogenic electron tomography, we reveal evolutionarily conserved features shared by the invasive forms of *Plasmodium falciparum* and *Toxoplasma gondii*. By comparing resting *Toxoplasma* tachyzoites with those primed to invade we also gain new insight into the very first steps in invasion. For this work, we take an interdisciplinary approach, adopting a mixed-scale dense neural network that enables efficient and objective processing of the data. Combined, the results provide new information on how these important parasites accomplish the essential step of invasion.

## Introduction

The phylum Apicomplexa includes several of the most prevalent and important human eukaryotic pathogens, such as the malaria-causing *Plasmodium spp*., and *Toxoplasma gondii* that can cause severe neurological disease in the developing fetus and those who are immunocompromised [1–3]. These obligatory intracellular parasites have a highly polarized cell shape and enter a host cell by deploying a remarkable machine at their anterior end known as the apical complex (AC), which is highly conserved and entirely specific to the ~6000 species in the Apicomplexa phylum [4, 5]. Coordinate interaction of subcellular organelles and distinct elements of the cytoskeleton that comprise this extraordinary and dynamic machine allows active invasion by these single-celled eukaryotes into other (host) eukaryotic cells [6].

Active invasion by tachyzoites, the rapidly growing life stage of *Toxoplasma*, involves protrusion of a unique spiral of tubulin-based fibrils known as the conoid, a structure found only in the coccidian subgroup of apicomplexans [5,7,8]. During protrusion the conoid, as well as two preconoidal rings at its anterior end, move outward, away from the cell body. When protruded, these structures are anterior to the apical polar ring (APR), which functions as the anchoring site for the subpellicular microtubules (SPMTs) [9]. The SPMTs are arranged in a gently spiraling pattern and extend ~2/3 the length of the parasite to provide mechanical support [10].

During the initial stages of invasion, two distinct, apically located secretory organelles, the capsule-shaped micronemes and the club-shaped rhoptries, secrete their contents [6]. Micronemes deposit their contents onto the surface of the parasites and these proteins facilitate surface attachment of the parasite to the host cell surface [11]. Next, rhoptry proteins are injected by an unknown mechanism from the rhoptry neck into the host cytoplasm [12]. This cargo includes RON2 which then somehow integrates into the host plasma membrane and binds to a micronemal protein, AMA1, on the parasite surface. The result is a ring of tight interactions that migrates down the parasite as it invades into the host cell [13]. Following secretion from the rhoptry neck, the rhoptry bulb injects its contents into the host cell, including effector proteins that enable the parasites to grow within and co-opt the host cell. Two additional components of the AC that are poorly understood but are likely of prime importance, are a series of apical vesicles (AVs) and a pair of intra-conoid microtubules (IMTs), along which most of the AVs are aligned, and all within the conoid space. Previous structural studies have revealed that the most anterior AV sits between the tip of one or two rhoptries and the parasite’s apical plasma membrane [14–18]. This AV has been suggested to be a part of the docking machinery for rhoptries [14, 15, 19], through interactions with the plasma membrane-bound rhoptry secretory apparatus (RSA). Despite these recent advances, the exact means by which rhoptry secretion occurs and the precise role of the AVs in this process, remain unknown.

Cryogenic electron tomography (cryo-ET) is a powerful tool to study cellular architecture in situ without using chemical fixatives or metal stains [20]. This method can visualize membranes and protein-based cellular structures in a near-native, frozen, hydrated state but it can be severely limited by the thickness and the inherently low contrast of biological samples. Recently, cryo-ET has proven successful in revealing the 3D organization of the subcellular organelles in the AC of extracellular, non invading *Toxoplasma* [8, 15, 19]; however, due to the transient nature of host cell invasion and the limited throughput of cryo-ET, it has not yet been possible to use this method to capture the invasion process itself. As an alternative to observing parasites in direct interaction with host cells, one can observe the biological machinery of extracellular parasites under internal calcium flux induced by the addition of calcium ionophore. This stimulation, among other things, is known to prime the parasites for invasion by initiating protrusion of the conoid and microneme secretion [21–24].

As with most biological processes, there can be a substantial amount of cell-to-cell heterogeneity seen upon stimulation of *Toxoplasma* tachyzoites with calcium ionophore [23]. This is likely because each cell is in a unique physiological microstate and so quantifying significant differences between two conditions necessitates analyzing enough cells to distinguish what is the result of the manipulation (e.g., ionophore stimulation) versus what is normal cellular heterogeneity. Yet, generating and accurately annotating Cryo-ET tomograms is a labor-intensive process, limiting the number that can be readily analyzed. In addition, a significant bottleneck to moving from cryo-ET data to biological understanding is distilling the grayscale tomography reconstructions into feature-annotated reconstructions, a process that until recently was almost entirely manual. To enable our analyses, therefore, a method for high-throughput, automated annotation was needed and so we employed and customized a protocol for using a special type of neural network, namely, the Mixed Scale Dense (MSD) network [25], that is designed to require fewer training examples than other neural networks. By using the MSD network, we show that it is possible to achieve accurate segmentations across multiple tomograms and use them to analyze different parameters in an unbiased and quantitative matter. We report here the results from such an analysis including several new findings about the ways in which apical vesicles associate with other cellular entities upon ionophore stimulation and as a prelude to invasion.

## Results

Following stimulation with the calcium ionophore, A23187, the conoid of *Toxoplasma gondii* tachyzoites harvested from infected monolayers protrudes in an irreversible way [21] and micronemal proteins such as MIC2 are deposited onto the surface of the parasites where they can be cleaved by proteases [22]. If harvested into Endo buffer containing high potassium levels, intracellular calcium levels remain stable and invasion is prevented [26, 27]. Using a phase microscope, we assessed the percentage of extracellular parasites that had undergone conoid protrusion under “stimulating” (1 μM A23187 in Hanks Balanced Salt Solution (HBSS)) and “non-stimulating” conditions (Endo buffer). The results (Figure S1a, Supplementary Material) showed ~80% of the tachyzoites had protruded conoids after 0.5-2 minutes of ionophore-stimulation while only ~8% had protruded conoids even after 10 minutes incubation in non-stimulating conditions. To assess microneme secretion, a western blot was used to detect secreted MIC2 (Figure S1b, Supplementary Material). An already cleaved form of MIC2 was detected after 2 minutes of ionophore-stimulation at room temperature, while no cleaved MIC2 was detected when parasites were incubated in non-stimulating conditions. We, therefore, harvested the tachyzoites into either Endo buffer or HBSS supplemented with 1 μM A23187 for ~0.5-2 minutes prior to transfer to a lacey carbon grid and plunge-freezing followed by imaging using cryo-ET. A total of 40 tomograms of stimulated and 8 tomograms of unstimulated parasites were generated in this way and a representative tomogram from each condition is presented in Movie S1 and Movie S2, respectively. Virtual sections from these tomograms are presented in Fig. 1 together with a model of known components of the AC.

**Fig. 1.**
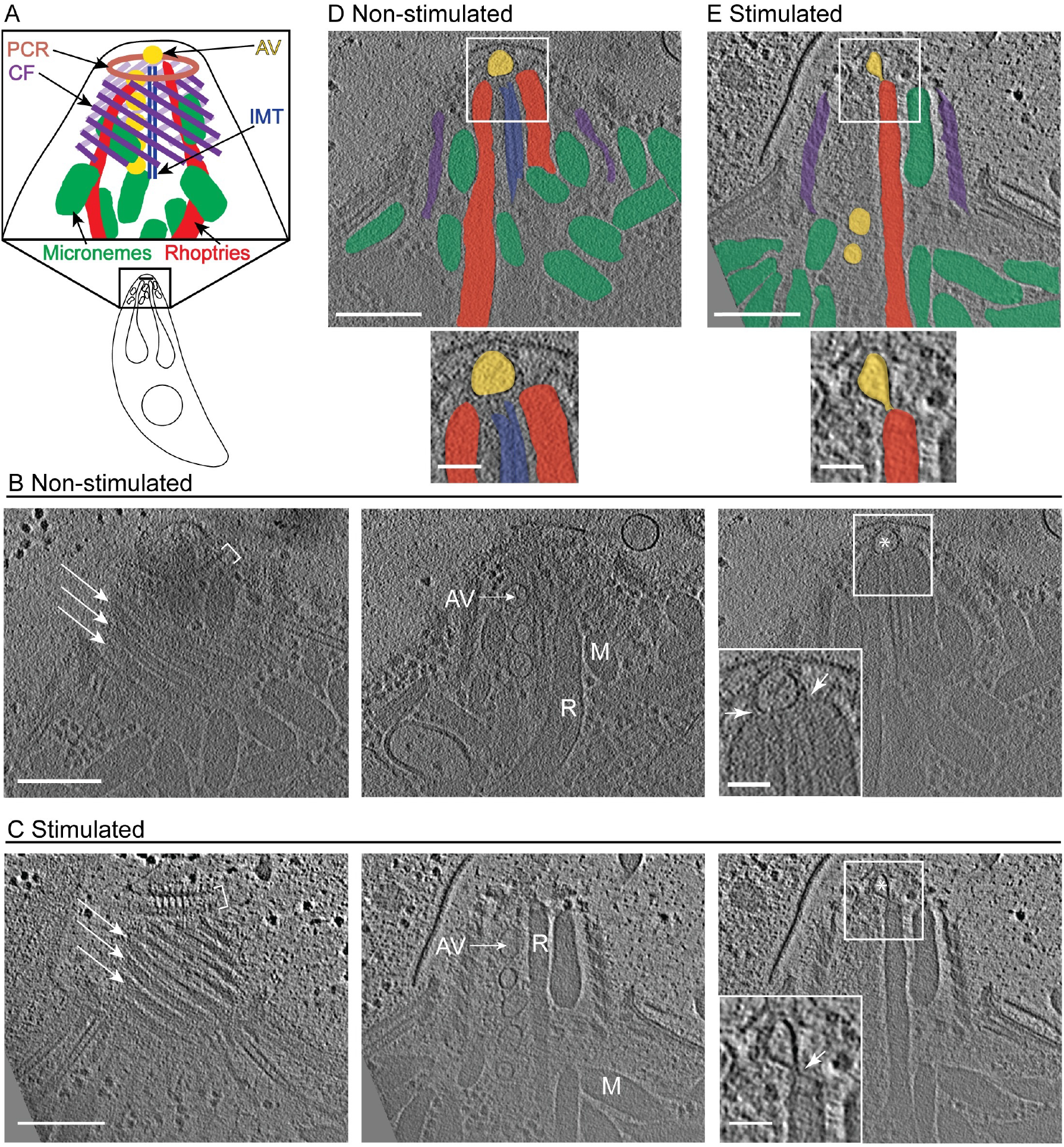
Cryo-ET reveals the organization of subcellular organelles in the apical complex upon ionophore-stimulation. **(A)** Cartoon of the *Toxoplasma* tachyzoite cell as well as an enlarged cartoon of the apical complex showing key subcellular structures: preconoidal rings (PCR; brown), conoid fibrils (CF; purple), intra-conoidal microtubules (IMT; blue), micronemes (green), rhoptries (red) and apical vesicles (AV; yellow). **(B)** Tomographic slices of a representative apical complex of non-stimulated tachyzoite showing the preconoidal rings (brackets), conoid fibrils (arrows), micronemes (M), rhoptries (R), IMT-associated AVs (AV), and the most anterior AV (asterisk). Scale bar, 200 nm. Inset is a zoomed-in view of the square showing the association (arrowheads) of the rhoptries with an AV. Scale bar, 50 nm. **(C)** as for (B) except the images are of a tachyzoite stimulated with ionophore. **(D)** and **(E)** are the slices shown in the far right of (B) and (C), respectively, with structures highlighted in colors as (A), and a zoomed in view of the square. Movies with the complete tomograms are available in S1 Movie (non-stimulated) and S2 Movie (ionophore-stimulated).

In all the tomograms, micronemes and rhoptries were clearly visible inside the space delimited by the conoid spiral. Microneme and rhoptry secretion are essential for successful invasion of apicomplexan parasites. Although it is hypothesized that both secretion events require a membrane fusion, no such interaction or fusion of the micronemes with the parasite membrane has been reported or was observed here. With regard to rhoptries, *Toxoplasma* carries up to a dozen such organelles, but only two rhoptry necks were observed within the intraconoid space of non-stimulated parasites. In stimulated parasites, the number of rhoptries that occupy the intraconoid space varied with 23 such cells displaying two rhoptries in the conoid space, 12 cells displaying only one rhoptry, and 5 that did not display any rhoptries in this space (58%, 30% and 12%, respectively, N=40). A recent cryo-ET study of *Toxoplasma* tachyzoites harvested in a buffer that was not specifically reported as inducing or inhibiting conoid protrusion, described a constriction in the rhoptry neck, a phenotype that could only be detected by reconstructing the 3D volume of the rhoptries [19]. In tomograms where the rhoptries were visible, we observed a similar constriction in the rhoptry neck in a majority of the stimulated parasites (23 of 36 cells) and also in 1 of 7 non-stimulated (Figure S2a, and Movie S3, Supplementary Material).

In addition to the micronemes and rhoptries, two IMTs and a series of up to 6 AVs were also seen in the intraconoid space. Relative to the plane of the electron microscopy grid, the IMTs were positioned in the center of the conoid in 13% of the cells (N=47); otherwise, these IMTs were positioned to one or other side of the intraconoid space (87%), as viewed from above. Intriguingly, there was a strong relationship between the position of the AVs relative to the IMTs and the position of the IMTs in the conoid space, such that the vesicles were almost always (31/33) found on the side of the IMT closest to the conoid fibrils (*φ*=0.87, Pearson correlation for two binary variables). When two rhoptries were found within the conoid space, they were always positioned on opposite sides of the IMTs, whereas when only one rhoptry was present (N=12), it was usually (10 of 12 parasites) on the side not occupied by the AVs (*φ*=-0.63, Pearson correlation for two binary variables). Tomograms of non-stimulated parasites revealed a single AV positioned at the apex of the cell, anterior to the IMTs, while tomograms of stimulated parasites revealed one, two, or even three AVs in this location (37, 3, and 1 such cells, respectively) (Figure S2b/c, Supplementary Material). Of greatest interest, however, was that in many of the ionophore-stimulated *Toxoplasma* parasites there was an intimate association between one or two rhoptries and the most anterior AV, including an apparent deformation of the AV membrane towards the rhoptry tip (Fig. 1c/e and Figure S2d/e, Supplementary Material). This point is further explored below.

To investigate whether the positioning of an AV between a rhoptry tip and the rosette is an evolutionarily conserved phenomenon, we repeated our analyses with merozoites of *Plasmodium falciparum*. These organisms are much more fragile than *Toxoplasma* tachyzoites and do not possess an obvious conoid structure and so we analyzed these only under non-inducing (no ionophore) conditions. The results revealed a single AV underneath the rosette (Fig. 2b) suggesting a unique and conserved role for the AV in rhoptry secretion in apicomplexan parasites.

**Fig. 2.**
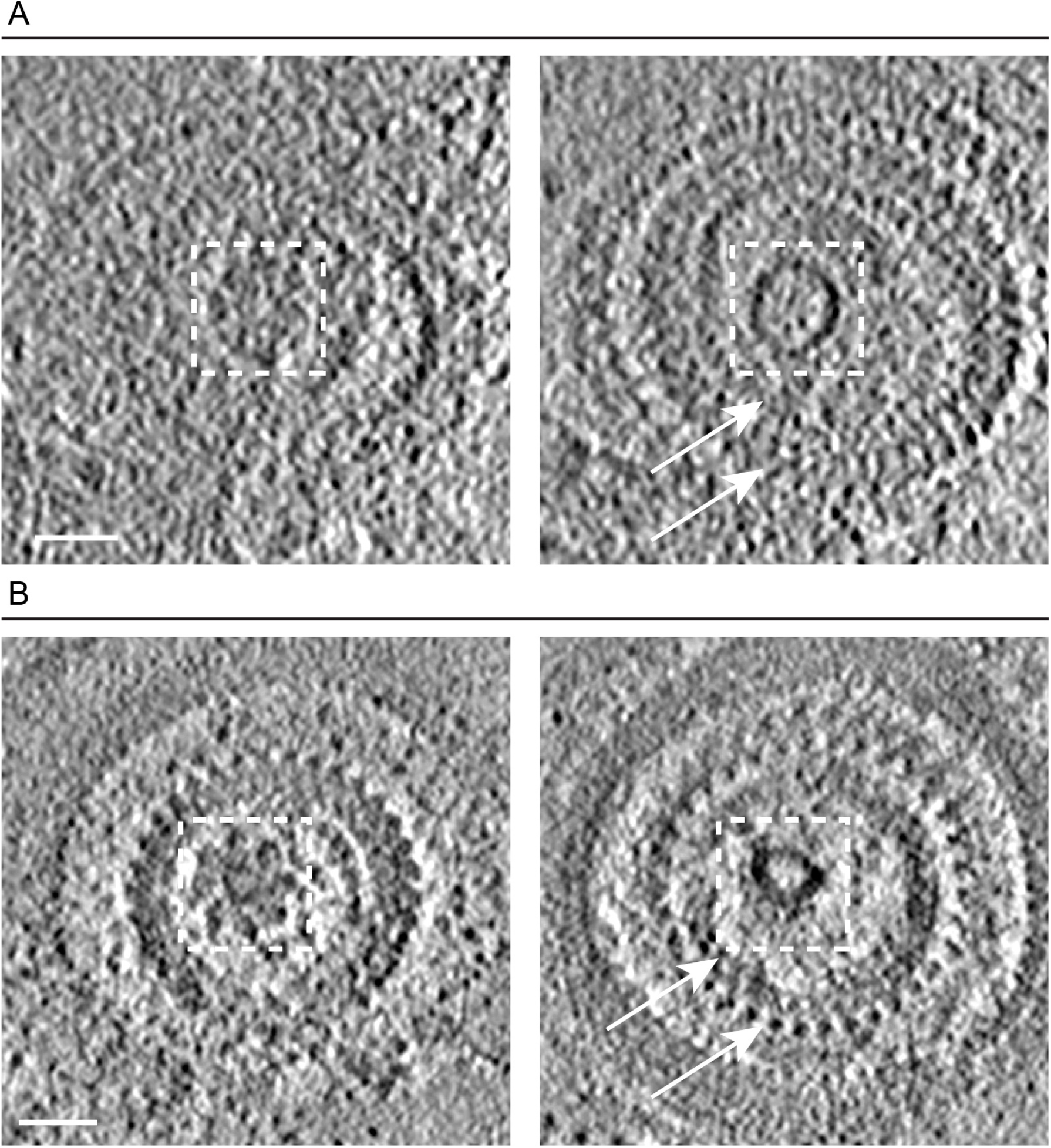
An apical vesicle is located underneath the rosette in *Toxoplasma gondii* tachyzoites and *Plasmodium falciparum* merozoites. **(A)** Left panel is a tomographic slice showing a top view of the rosette of a *Toxoplasma* tachyzoite. Right panel is a different z-slice from the same tomogram showing the anterior apical vesicle (AV) and the preconoidal rings (arrows). The dashed squares mark the same area in the tomographic slice showing that the AV is positioned about 25 nm underneath the rosette. **(B)** As for (A) except the tomogram showing the rosette, AV, and two (out of three) polar rings of a *Plasmodium falciparum* merozoite. Scale bar, 50 nm.

### A mixed-scale dense neural network allows fast and accurate annotation of distinct segments using a limited amount of training data

To be confident that possibly subtle differences observed under two or more sample conditions are significant, large amounts of data are needed. Related to this, one of the challenges of using cryo-ET data for studying biological phenomena has been the very large amount of time needed to annotate and then analyze the tomograms; hence the need for an automated approach for image analysis. Traditional neural networks for image analysis, however, typically require vast amounts of “training data,” produced by manual annotation, making them difficult to employ in practice. To meet this challenge, we adopted and customized the Mixed-Scale Dense (MSD) Neural Networks (NN) approach [25] to perform image segmentation and classification. Unlike traditional neural networks, such as U-Net and SegNet [28, 29], MSD networks are specifically designed for scientific imaging data where there is limited amount of ground-truth training data. These networks operate by removing the upscaling and downscaling in typical networks and replacing these operations by mathematical stencil operators which perform the convolutions across multiple scales (“mixed scale”) without requiring generation of intermediate objects that can only communicate with nearby layers. Consequently, all layers can be connected together, hence the idea of a “dense” network.

Here, we adopt these MSD networks to quantify the effect of ionophore-stimulation on the organization of the AVs within the AC. For this application, we used the following iterative process. The initial training data included manual annotation of just 3-4 adjacent slices from 7 reconstructed tomograms. The slices were chosen by their relevance to the biological question and therefore all included one or more AVs. All the relevant features (AVs, rhoptries and IMTs) in each slice were annotated. Both the IMTs and rhoptries were challenging for the network to annotate correctly due to their limited number per tomogram, at most only two of each, and their similarity to other, more abundant, structures namely the SPMTs and the micronemes, which in cross section can look quite similar to a rhoptry neck in a single slice. To overcome this limitation, one label was used for both the SPMTs and IMTs and another for the rhoptries and micronemes. Combining related structures into a single label reduced errors in the network output and in most cases the difference between the two combined entities could be readily discerned in the NN output (e.g., the location of the SPMTs and IMTs are completely different, a feature that the NN did not pick up on as readily as other differences from the other structures present; they also differ substantially when viewed in three dimensions which the NN was not able to assess) (Figure S3a/b, Supplementary Material).

Rather than apply this initially trained network to all of the acquired tomograms, we used an iterative bootstrap procedure: we applied the network to 3-6 biologically relevant, adjacent slices from 11 additional tomograms, and then manually fine-tuned the results to create an expanded training set. Making these corrections was much faster than manual annotation de novo and allowed for 43 additional annotated slices to be used for training, which led to a considerable increase in accuracy when the network was applied to all tomography data.

To characterize and assess the NN derived annotations relative to ones made by expert human annotators, we compared the level of agreement over two parameters, pixel-wise error and connected-component error. The percentage agreement between two expert human annotators for the AVs and the micronemes/rhoptries was between ~80-100% for the two parameters, respectively; this level of agreement was similar to that between each of the human experts and the NN (Figure S3c, Supplementary Material). For the microtubule’s annotation, the pixel-wise agreement was ~70% for the two human annotators, making this the structure for which the human annotators had the least agreement, probably due to the short subpellicular microtubules seen in crosssection. This could make it challenging for the NN to learn as reflected in the ~50% agreement between the human and NN annotations, although due to the large standard deviation, the difference between this and the ~70% agreement for the two human annotators was not statistically significant. This iterative training based on manual correction improved the accuracy of the NN in annotating prominent structures, as well as in detecting low-abundance features (e.g., AVs; Figure S4, Supplementary Material), yielding a highly accurate annotation for tomograms that were not used in the training data (Fig. 3). Thus, the NN process enables faster and more objective analyses of the data, including characterization of subcellular components and their possible interaction under stimulating and non-stimulating conditions, as described further below.

### Exposure to a calcium ionophore promotes an intimate interaction between the most anterior AV and rhoptry tips

To quantify the precise changes induced by calcium ionophore, we used the NN to analyze multiple tomograms of ionophore-stimulated and non-stimulated extracellular parasites. These changes led us to explore the possibility that stimulation is affecting the position of the most anterior AV relative to the IMTs by measuring the distance between these two structures. The results showed similar distances between the AV center-of-mass (COM) and the IMT for stimulated (median=79.3 nm, mean ± SD =79.3±14.6 nm, N=31) and non-stimulated (median=73 nm, mean ± SD =72±15 nm, N=8) parasites (Fig. 4c). We also measured the distances between the AVs’ boundaries (membrane) and the IMT (Fig. 4d), again revealing a similar distribution of distances (median=53.5 nm, mean ± SD =53.3±13.7 nm, N=31 for stimulated parasites and median=48.4 nm, mean ± SD =47.1±19 nm, N=8 for non-stimulated parasites), meaning the distance is not affected by the ionophore stimulation. Lastly, we measured the angle between the AV and a best fit line through the nearest IMT and observed that this did not change upon stimulation, as presented in Fig. 4e (median=18°, mean ± SD =19.6±11.7°, N=31 for stimulated parasites and median=15.6°, mean ± SD =18.8±15.4°, N=8 for non-stimulated parasites).

**Fig. 3.**
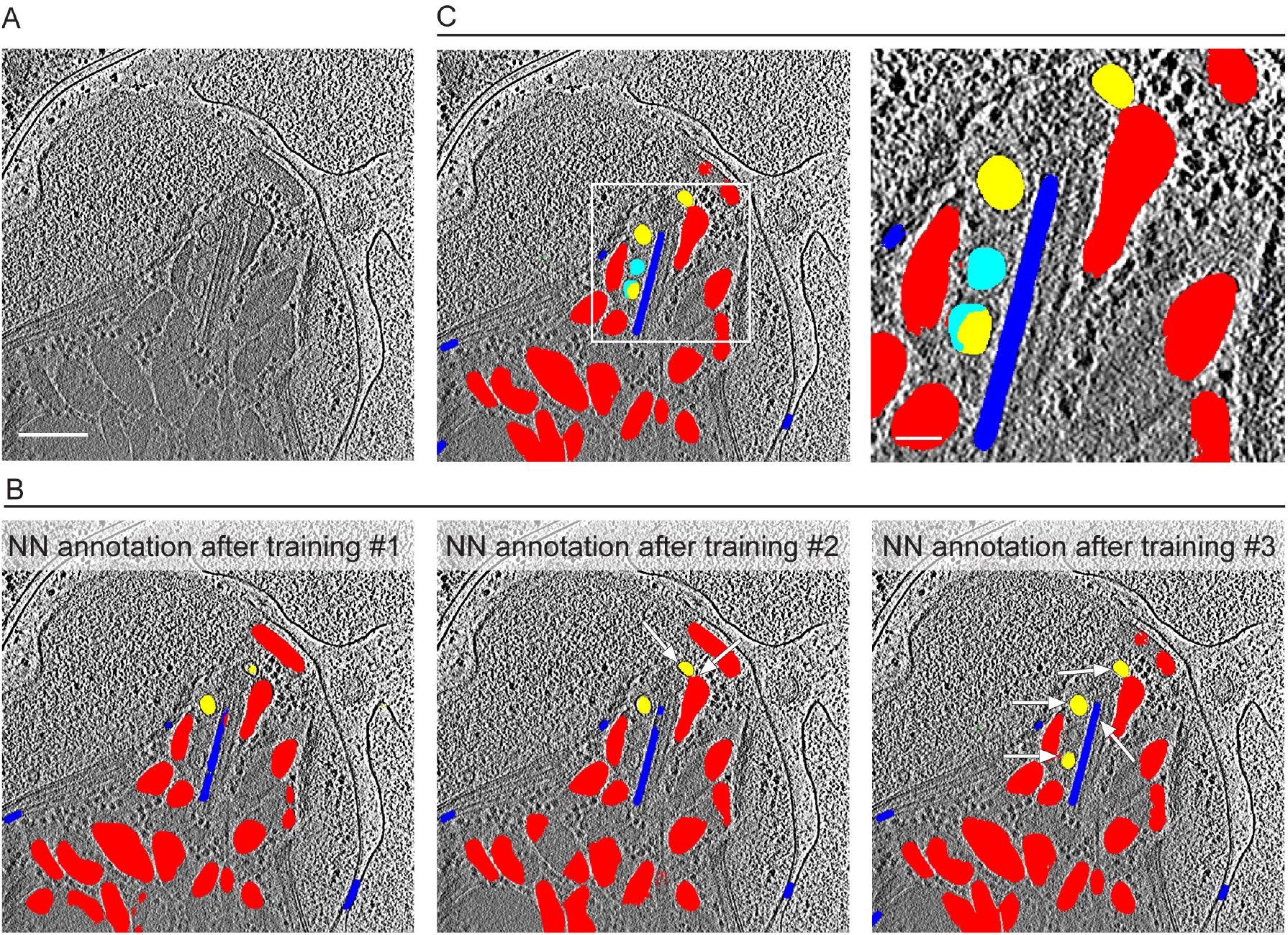
Annotation of distinct elements in the apical complex of *Toxoplasma* tachyzoites using a mixed-scale dense neural network. **(A)** A representative slice from a tomogram that was not used in the training data and that is shown annotated in (B) and (C). Scale bar, 200 nm. **(B)** Annotation produced by the neural network after different iterative rounds of training (see results section for details) overlaid on the tomographic slice from (A) showing the AVs (yellow), rhoptries and micronemes (red), and microtubules (blue). Note the small but marked improvement in the annotation accuracy of the AVs, rhoptry, and IMT with each additional training as pointed by the arrows. **(C)** Manual correction (cyan) of the NN annotation after training #3. A zoomed-in view of the square in the left panel is shown in the right panel. Scale bar for the zoomed-in view, 50 nm.

**Fig. 4.**
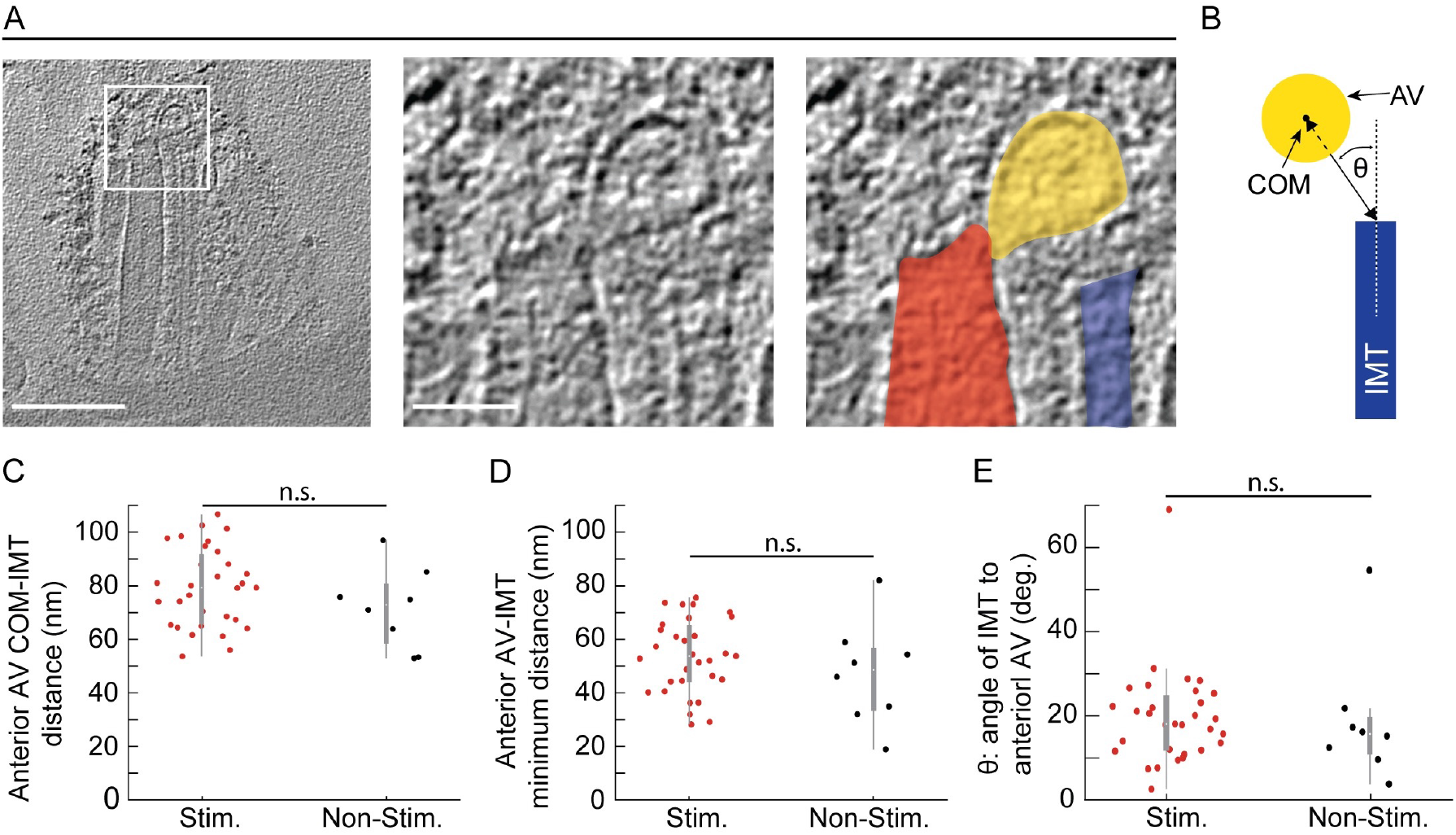
The positioning of the anterior AV relative to the IMT in *Toxoplasma* tachyzoites does not change upon ionophore-stimulation. **(A)** Left panel: a tomographic slice of the AC of an ionophore-stimulated tachyzoite. Scale bar, 200 nm. Middle panel: a zoomed-in view of the square in the left panel. Scale bar, 50 nm. Right panel: as for the middle panel image but highlighting in color the tip of the rhoptry (red), anterior AV (yellow), and IMT (blue). **(B)** A schematic showing the parameters used to assess the relative positions of the IMT (blue) and anterior AV (yellow). **(C)** The distance from the anterior AV’s center-of-mass (COM) to the tip of the IMT in stimulated and non-stimulated tachyzoites. Each dot is the result for one tomogram. **(D)** As for (C) except the distance from the IMT tip to the closest part of the anterior AV’s membrane is shown. **(E)** The angle (θ) subtended by the long axis of the closest IMT and a line connecting the COM of the AV to the IMT tip. There was no statistically significant difference (“n.s.”) between the stimulated and non-stimulated parasites for any of these three comparisons.

One of the most striking changes we observed upon ionophore-stimulation was the deformation of the anterior-most AV in the form of a protrusion directed toward the tip of a nearby rhoptry. We observed this deformation in 29 (81%) of the stimulated parasites (N=36), whereas it was not observed in any unstimulated parasites (z=4.162, 99.9% for Two-Sample Z-test for proportions). This deformation could be quantified using the output of the NN annotation to determine the membrane curvature of each vesicle. This curvature was assessed in the 2-D virtual slices that were within ~8 nm of each vesicle’s center of mass and was averaged as a function of angle around the vesicle (Figure S5, Supplementary Material). The results were plotted in 10 degree increments, starting at the voxel of the vesicle that is closest to a voxel of the interacting rhoptry (Fig. 5a/b). In stimulated parasites, this analysis showed a clear transition from high to low curvature when moving away from the point of interaction (i.e., a teardrop-like shape). This effect was absent in non-stimulated parasites where the curvature was relatively constant, reflecting their near spherical shape and similar to the shape of the more posterior, IMT-aligned AVs where a similar quantification was performed but the values were aligned relative to the vesicle voxel that was closest to an IMT voxel.

**Fig. 5.**
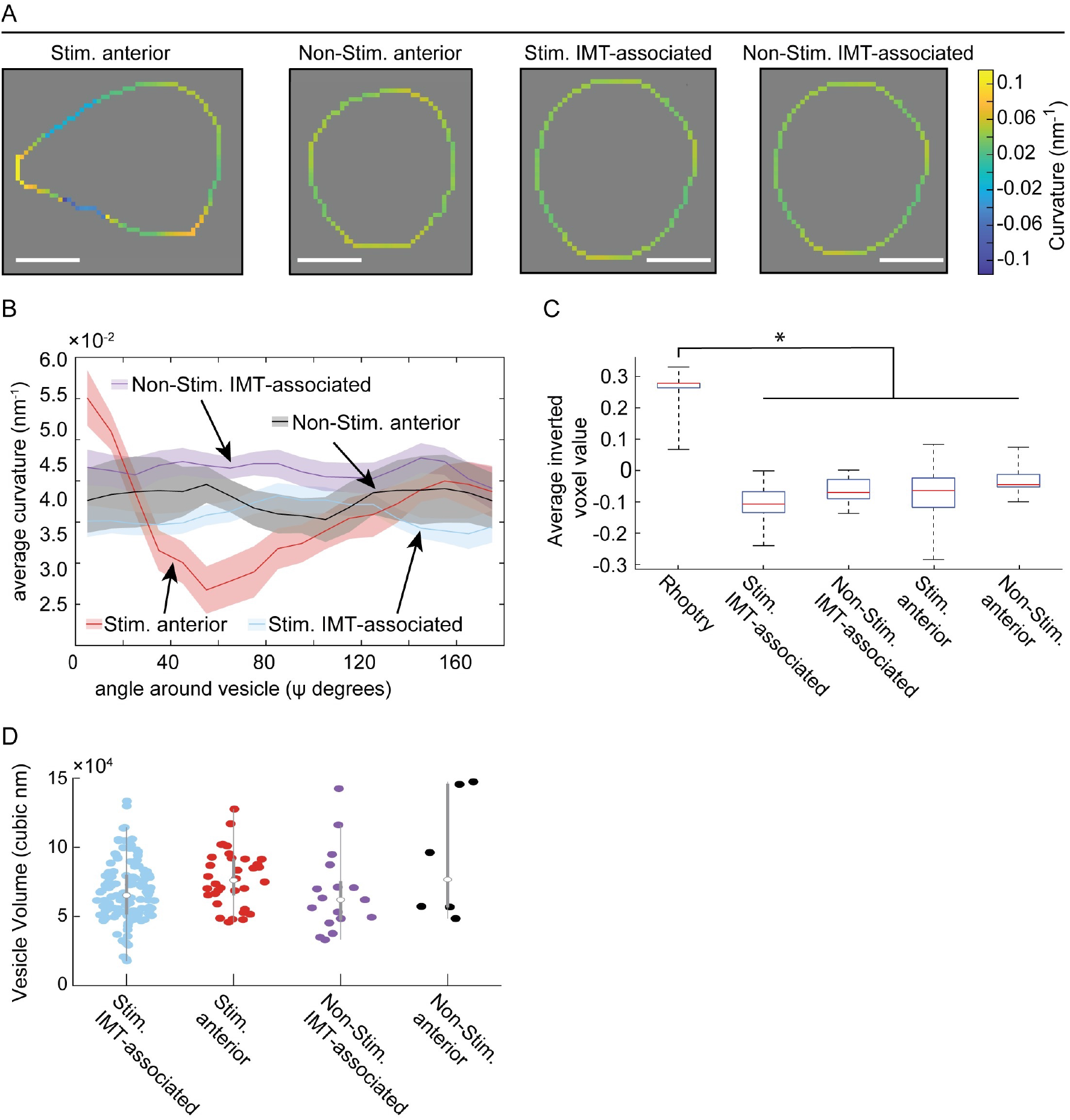
Ionophore-stimulation of tachyzoites results in a distortion of the anterior AV’s shape but not its density relative to the other AVs. **(A)** The curvature of representative anterior and IMT-associated AVs in tachyzoites. The curvature around the vesicles was measured for each pixel on the periphery of the vesicle in xy slices. See Materials and methods for more details. Vesicles were aligned by their shortest distance between each vesicle’s COM and either the rhoptry tip or the IMT, for anterior and IMT-associated vesicles, respectively (see details in Figure S5, Supplementary Material). The heat-map reflects the positive and negative curvature as “warmer” and “cooler” colors, respectively. Scale bar 20 nm. The three-dimensional (3-D) annotation of the AVs are presented in S1 Movie (non-stimulated) and S2 Movie and S4 Movie (stimulated). Note that the tear drop shape was also apparent in 3-D. **(B)** The averaged curvature for all AVs in stimulated (Stim.) and non-stimulated (Non-Stim.) tachyzoites, measured every 10 degrees as in (A). **(C)** The density of the rhoptries (mean ± SD =-0.259±0.054, N=35), and anterior and IMT-associated AVs under stimulated (Stim.; anterior mean ± SD =0.082±0.083, N=34, and IMT associated mean ± SD =0.103±0.048, N=119) and non-stimulated (Non-Stim.; anterior mean ± SD =0.03±0.058, N=6, and IMT associated mean ± SD =0.656±0.042, N=17) conditions. The only statistically significant difference (p<0.05) was between the rhoptry and the AVs in each of the four conditions. **(D)** The volumes of the anterior and IMT-associated AVs under stimulated (Stim.; anterior mean ± SD =7.7±2 X10^4^ nm^3^, N=34, and IMT-associated mean ± SD =6.8±2.2 X10^4^ nm^3^, N=119) and non-stimulated (Non-Stim.; anterior mean ± SD =9.2±4.5 X10^4^ nm^3^, N=6, and IMT-associated mean ± SD =6.7±2.9 X10^4^ nm^3^, N=17) conditions. None of the pair-wise comparisons showed a statistically significant difference.

The teardrop shape of the rhoptry-interacting AV in stimulated parasites suggests either a fission or a fusion event was taking place. Examination of the region where the two structures are closely apposed in greater detail revealed an apparent continuity between the lumens of the rhoptry and interacting AV in 12 out of 36 tomograms of ionophore-stimulated parasites (representative images are shown in Fig. 1b and Fig. 4a). To distinguish between fusion and fission processes, we compared the density of the AV’s contents to that of the rhoptries’; fission might predict that they should have similar contents (since the interacting AV would be very recently derived from rhoptries), whereas if they are of completely different origins and are just initiating fusion, they might have very different cargo, as reflected in the density of their contents in the tomograms. The results (Fig. 5c) showed that the median density of the AVs was similar, regardless of whether they were interacting with the rhoptry or one of the more posteriorly located, IMT-associated vesicles, and that this density was substantially less than the rhoptries. These results suggest, but certainly do not prove, that the AVs have a distinct cellular origin from rhoptries and that the interacting AV is in the process of fusion with, rather than budding from, the rhoptry tip. We also compared the volumes of the AVs. The results (Fig. 5d) showed that the volumes were similar between the AVs, regardless of their position or whether they are stimulated. This does not distinguish between the fusion and fission hypotheses but does further indicate that the two structures are distinct and that their intimate interaction is a stable state because a significant volume has neither flowed into nor out of the interacting AV.

### The AVs associate with the IMT and with each other

In addition to the AV positioned anterior to the IMT, we observed up to 6 AVs aligned along the IMTs in the intraconoid space, with a mode of 3, regardless of whether they had been exposed to calcium ionophore (Fig. 6a). These AVs were usually lined up with a regular spacing between them and a uniform distance from the IMTs, suggesting an association between these various components (Fig. 6b). In 7 tomograms (5 stimulated and 2 non-stimulated parasites), we also identified apparent AVs posterior to the IMTs (Figure S6a, Supplementary Material). Closer inspection of the IMT-aligned AVs revealed a density partially surrounding the vesicle in the majority (75%) of the tomograms, with a similar density also observed surrounding the AVs positioned anterior to the IMTs (Figure S7a-d, Supplementary Material). Using the tomographic data to examine this structure more carefully revealed that in at least 31 out of 48 tomograms the density partially coats the AVs, and in 10 instances it resembled the rosette that was previously characterized as part of the RSA found between the rhoptry-interacting AV and the parasite’s plasma membrane [19] (Figure S8a-d, Supplementary Material). The difference in our data from those previously reported is that we observe these rosette-like structures associated with more than one AV in a single AC and even around AVs that are IMT-associated, not just those that are anterior-most or interacting with a rhoptry tip.

**Fig. 6.**
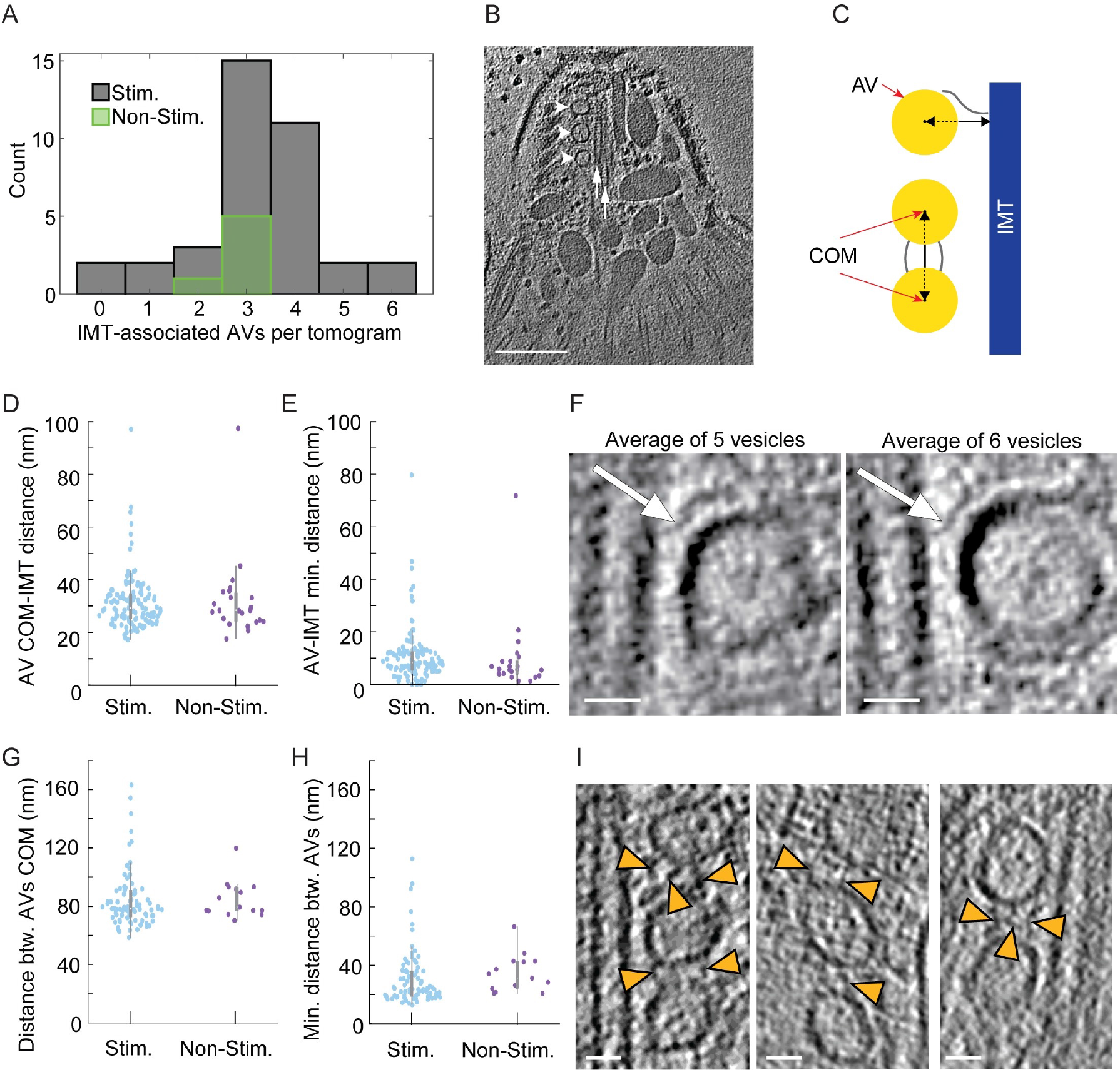
The AVs interact with the IMTs and with each other. **(A)** The number of IMT-associated AVs in the AC of ionophore-stimulated (Stim.) and non-stimulated (Non-Stim.) tachyzoites. **(B)** A tomographic slice of the AC of a stimulated tachyzoite showing three AVs (arrowheads) associated with the IMTs (arrows). Scale bar, 200 nm. **(C)** A schematic showing the parameters used to assess the distance between the AVs (yellow) and between each AV and the IMT (blue). Measurements began at either the AVs’ COM (red arrows; middle and bottom AVs) or their delimiting membrane (red arrow; top AV). **(D)** The distance from each AV’s COM to the IMT in stimulated and non-stimulated tachyzoites. Each dot is the result for one AV. **(E)** As for (D) except the distance from the AV membrane to the IMT is plotted. **(F)** Subtomogram averaging of all IMT-associated AVs from two separate tomograms of stimulated tachyzoites. Left panel is the output from averaging 5 IMT-associated AVs from one tachyzoite; right panel is the result from averaging 6 IMT-associated AVs from a different tachyzoite. AVs were aligned by the minimum distance between the AVs and the IMT. This averaging reveals an apparent density between the two structures (arrows). Scale bar, 20 nm. **(G)** The distance between neighboring AV’s COM in stimulated and non-stimulated tachyzoites. **(H)** As for (G) except the distance between neighboring AV’s delimiting membrane is plotted. **(I)** Tomographic slices from 3 ionophore-stimulated tachyzoites showing a density (arrowheads) reproducibly seen between the IMT-associated AVs. Scale bar, 2O nm. No statistically significant difference was seen for any of the pairwise comparisons in the five datasets shown (parts (A), (D), (E), (G) and (H)).

The fact that the AVs are generally aligned on just one side of the IMT pair raises the question of how the two IMTs might differ from each other. Our results showed that within any given tomogram, regardless of whether the parasites had been exposed to calcium ionophore or not, the two IMTs of any given pair were similar in length (Figure S6b, Supplementary Material). We also observed an internal striation density within the lumen of both IMTs, but no external densities regularly coating one IMT or another, leaving unanswered what determines the side of the IMT with which AVs will associate.

Next, to determine if ionophore stimulation alters the IMT pair overall, we examined the stimulated and non-stimulated parasites and found that both the length (350±60 nm for stimulated, and ~300±30 nm for non-stimulated tachyzoites) and curvature (~0.7X10^-4^ nm^-1^ for stimulated, and ~1X10^-4^ nm^-1^ for non-stimulated tachyzoites) did not significantly change upon stimulation (Figure S6b/c, Supplementary Material).

To determine if ionophore-stimulation caused other changes in the AV/IMT interaction, we measured the median distance between each AV’s center of mass (COM) and the nearest IMT. The results (Fig. 6d) showed no change upon stimulation (median= 29.2 nm, mean ± SD =31.7±10.8 nm, N=119 for stimulated parasites and median= 28.4 nm, mean ± SD =32.1±15.6 nm, N=23 for non-stimulated parasites). Similarly, the distance measured between the AVs’ boundaries and the IMTs showed a tight distribution that did not change upon stimulation (Fig. 6e). Given their approximately spherical shape, this agreement between the results when measuring from the COM and outer edge of the AVs also indicates that the AVs are of similar size in the two conditions. To explore possible interactions that can maintain the regular distance between the vesicles and an IMT, we extracted the densities in the interface between the two structures and aligned them according to the minimal distance vector between the AV’s boundary and the IMT. Two subtomogram averages, each of which was created from a different tachyzoite (one that included 5 vesicles and the other 6 vesicles) revealed a similar ~25 nm long density apparently connecting the vesicle and the nearest IMT (Fig. 5f). We also measured the distance between the COM of neighboring vesicles to determine if ionophore-stimulation caused changes in the spacing between them. The results (Fig. 6g) showed no significant difference upon stimulation (median= 78.4 nm, mean ± SD =84.1±19.3 nm, N=84 for stimulated parasites and median= 79.4 nm, mean ± SD =85±12.6 nm, N=15 for non-stimulated parasite). Similarly, the distance between the AVs’ boundaries (Fig. 6h) did not change significantly upon stimulation (stimulated, mean ± SD =30.6±18.5 nm median=23.6 nm, N=84; and non-stimulated, mean ± SD =34.2±12.7 nm median=31.2 nm, N=15). To determine if the distribution of the AVs along the IMT change upon ionophore stimulation, we measured the distance between the most anterior IMT-associated AV to the tip of the IMT. The results showed no change upon stimulation with a mean of ~40±10 nm for both stimulating and non-stimulating conditions (Figure S6e, Supplementary Material). Interestingly, we frequently detected 1-3 densities apparently connecting neighboring vesicles, but we were not able to average the densities due to structural heterogeneity (Fig. 6i). The average length of the intervesicular connections, as determined by manual annotation of 43 filaments, was 28±5 nm while the average distance between the AVs at their closest points was 22±4 nm. Vesicles that did not have any detectable density connecting them appeared more distant from each other with average distances of 45±11 nm (Figure S6f/g, Supplementary Material). These data suggest the existence of distinct structures that both tether the AVs to the IMTs and that maintain a constant distance between IMT-associating AVs.

## Discussion

Cryo-ET enables the observation of intact cells and cell fragments with nanometer scale 3D resolution, thereby revealing features that otherwise might be missed in analysis of electron micrographs using thin sections. Cryo-ET also benefits from involving no chemical fixation, only flash-freezing of samples, preserving the near-native cellular state and avoiding fixation artifacts. Despite these merits of cryo-ET and cryogenic sample preparation, in order to draw reliable conclusions many samples must be analyzed and careful annotation of the tomograms is necessary to quantify observed features. This manually intensive process of annotation remains a bottleneck of the cryo-ET workflow. Here, we describe how we tailored the Mixed-Scale Dense Neural Network to answer a biological question while training on very little manually annotated data. The resulting network provided a robust, automated workflow for image-segmentation-and-recognition across a large number of results. While the NN learning process is not controlled by the user, our protocol for generating the training data followed four key aspects that improved its annotation. (1) We annotated a few slices from different tomograms; (2) We carefully picked the slices used for training to include the most information on our features of interest; (3) We annotated full slices with multiple labels to include information on the relative location of the features; (4) We annotated adjacent slices to reflect the consistency in the shape and size of the features in 3-D. The resulting network could be applied across multiple tomograms from different samples and enabled us to obtain accurate annotations from slices that were not part of the training set. Importantly, the same network could be applied to both the biological conditions being examined blindly, avoiding bias that might be introduced by manual annotators alone, and the NN-generated annotations were generally as similar to those of two expert annotators as those two annotators were to each other. We were also able to overcome the challenge presented by differences in the signal-to-noise ratios in different tomograms as the NN successfully annotated tomograms that were not part of the training set. As a result, we were able to quickly and accurately annotate and then quantify features in a large number of tomograms. Subtomogram-averaging of the features thereby detected allowed us to discover structures not previously reported (e.g., densities connecting AVs to the IMTs), as well as identify differences produced by ionophore stimulation (e.g., direct interconnection of the lumens of the anterior-most AV and a rhoptry tip). Using corrected NN annotations as additional training data to improve the NN’s accuracy is an important advance over existing pipelines as it reduces the manual annotation time dramatically, making a major step forward in tackling biological problems in which extensive, laborious manual segmentation and annotation is a significant bottleneck in extracting information from data.

Inspection of the tomograms revealed that the plasma membrane of the tachyzoites was clearly visible and intact in less than half the tomograms under both inducing and non-inducing conditions. We note that tomograms presenting disrupted plasma membrane (likely due to the freeze plunge step to remove excess liquid by blotting) could have added noise to the relative positions of the subcellular organelles, but we observed no consistent difference between tomograms of parasites with and without visibly intact plasma membranes. Micronemes were always seen occupying the conoid space suggesting that secretion takes place at the tip of the parasite, beyond the conoid, as the region where the plasma membrane can be most easily accessed by such a large structure [22]. Microneme secretion was previously shown to involve ferlin 1 (TgFER1) and TgDOC2, both C2-domain-containing proteins [30, 31]. C2-domains conditionally engage with lipids in a calcium-dependent manner, facilitating membrane fusion [32]. It has therefore been suggested that micronemes directly fuse with the parasite membrane to discharge their contents [11, 30]. The calcium ionophore used here was previously shown to induce microneme secretion at 37°C [22]. Here, we used cooler, room-temperature conditions and still observed such a release. Although we hoped to possibly capture active microneme secretion in our tomograms, we did not identify any apparent fusion between these organelles and the plasma membrane. Part of the reason for this could be that the plasma membrane was clearly visible and generally intact in only ~52% of the tomograms under both conditions (25 out of 48 tomograms). It may also be that the fusion process is very short-lived and therefore difficult to capture.

Rhoptry secretion has been detected only upon contact with a host cell, but the exact nature of the signal required and the process leading to such secretion are unclear. Docking of rhoptries to the anterior AV within the conoid prior to egress could potentially enable a very rapid response once contact with a host cell has been made. Constriction of the rhoptry neck was recently reported in *Toxoplasma* tachyzoites, in areas of the neck that are devoid of luminal filaments [19] and our results indicate that this phenomenon is not stimulation-dependent. The purpose of these constrictions and whether the arrangements of the luminal filaments are dynamic remain to be determined.

Within the superphylum Alveolate, two phyla related to Apicomplexa are Dinoflagellata and Ciliata. In the ciliate *Paramecium*, the regulated secretion from the secretory organelles, trychocysts, involves docking to the plasma membrane at exocytosis sites where intramembranous particles form a rosette [33]. In Apicomplexa, an AV is sandwiched between the rosette and the rhoptries and it was recently suggested that these AVs facilitate the docking of a rhoptry to the parasite membrane through a newly described entity named the rhoptry secretory apparatus (RSA) [19]. The role of the AVs in rhoptry secretion has long been speculated [14, 34], and they were previously described in *Toxoplasma, Cryptosporidium, Eimeria, Sarcocystis, and Besnoitia* [15–17, 19, 35]; here, we extend their detection to *Plasmodium*. We observed that upon treatment of *Toxoplasma* tachyzoites with a calcium-ionophore, a commonly used stimulator of egress and invasion, a distended AV was seen fused with a rhoptry tip, and this connection appears to be stable enough to be captured in most such tomograms. This result is in agreement with another observation that links calcium signaling to rhoptry secretion, where depletion of the calcium-sensitive ferlin 2 (TgFER2) directly blocks rhoptry secretion [36]. Moreover, mass spectrometry analysis identified TgFER2 as an interacting partner of Nd9 together with Nd6, both of which are part of a conserved complex essential for rhoptry secretion; Nd6 was localized to the apical tip, presumably where the anterior AV is located [15]. Whether TgFER2 facilitates the AV-rhoptry fusion remains to be determined.

The difference between the density measured for the AVs and the rhoptry favors a model where these two entities have distinct origins, rather than AVs being derived by fission from rhoptries, unless a mechanism exists to pass into the AV only a subset of the rhoptry contents. It is possible that the AV/rhoptry interaction stabilizes the position of the latter in the intraconoid space, keeping it docked through the RSA to the plasma membrane as the conoid protrudes and retracts and micronemes enter the conoid space on their way to secrete. Assuming that rhoptry proteins are indeed transported to the host cell through the RSA, it is also possible that the interaction reflects a given rhoptry’s commitment towards secretion. It is unclear if more than one rhoptry secretes its contents during a tachyzoite’s invasion of one host cell. Interestingly, although unstimulated tachyzoites invariably displayed two rhoptries within the intraconoid space, upon stimulation we mostly observed interaction between an AV and only one rhoptry tip, even when two rhoptries were in close proximity. The apparent fusion of an AV to the rhoptry tip that we see upon ionophore-stimulation is similar to what was reported for Cryptosporidium sporozoites, where a continuity between the AV and the single rhoptry was observed, albeit without any stimulation [19].

The size of the AVs and the average distance between neighboring IMT-aligned AVs predict that a single IMT could interact with up to 6 vesicles, as we also observed, although we observed a mode of just 3. It is unclear if there is recruitment and transport of additional AVs to the IMTs but the appearance of vesicles with all the properties of an AV, posterior to and not associating with the IMTs suggests that the recruitment of AVs and their association with the IMT is a dynamic process. Whether there is, in fact, a dynamic equilibrium of AVs will require identification of their cargo and/or other markers that allow them to be specifically labeled.

The similarity between the IMT-aligned and rhoptry-associating AVs in volume, and density, and the appearance of more than one anterior, non-IMT-associating AV in the AC of stimulated parasites supports the previous suggestion [14, 19] that all AVs have a common origin. It was also suggested that the anterior AV-rosette association is independent of rhoptry docking [19]. We extend those results by reporting that the rosette structure that was previously seen within the RSA of the anterior-most AV is also apparent next to other AVs, either associated or not with the IMTs. *Toxoplasma* tachyzoites are capable of multiple invasion attempts where the rhoptries secrete their content into a host cell without completing invasion [37, 38]. To accomplish this, it could be that an AV stored along the IMT relocates to the apex of the cell where it can interact with a newly extended rhoptry and support its secretion after a failed invasion attempt involving a previous AV-rhoptry interaction. In the ciliate *Paramecium*, the intramembranous particles that form a rosette structure at the exocytosis site disappear following secretion [33]. Our data do not address whether the apical rosette observed here persists after secretion but the fact that we see such structures associating with more than just the anterior-most AV would allow for a new RSA to be transported to the apex of the parasite together with a new vesicle to enable multiple secretion events. Our results examining *Plasmodium falciparum* merozoites showed that association of a rosette with an anterior AV is conserved in that genus, although we did not examine sufficient numbers to address whether there are multiple AVs, IMTs or the other features seen in *Toxoplasma* tachyzoites. Likewise, we did not examine merozoites after ionophore-stimulation and so cannot address whether calcium signaling might play a similar role in these parasites as it does in *Toxoplasma*.

Transport of vesicles along the IMTs might be mediated in part by known motor proteins that are similar in size to the densities observed between the vesicles and the IMT, such as dynein or kinesin. Of note, however, the distance between the IMT to the anterior AV is greater than its distance to the IMT-associated AVs, meaning the connection between them must be altered for an IMT-associated AV to reposition as a new anterior AV.

Although no direct connection was detected between the IMTs or their associated AVs and the conoid fibrils, we did observe that the majority of the IMTs were not centered within the intraconoid space. We speculate that the asymmetric organization of features within the intraconoidal space might reflect the shape of the tachyzoite cell; i.e., the crescent-shape of the parasite could influence how it is lying on the surface of the grid during sample preparation, and potentially be enhanced by compression of the cells (due to blotting). Unfortunately, our limited field of view did not allow us to observe the entire tachyzoite or its overall curve, but it seems most likely that they would rest on one or other side since lying with the curve facing up or down would be much less stable. While we did note the presence of an apparent density or tether that was reproducibly present between the AVs and IMT, as well as between AVs aligned along this structure, we did not find any difference between the two IMTs that might explain which IMT interacts with the AVs. Ultimately, to uncover the exact means by which the AV supports rhoptry secretion, the structure and organization of the entire apical complex will need to be examined in parasites captured in the act of invasion of host cell, a technical challenge but within the scope of emerging technologies with cryo-ET after focused ion-beam milling [39], super-resolved cryogenic correlative light, and electron microscopy [40] and high-throughput tomogram data collection and processing.

## Materials and methods

### Parasite maintenance and cell culture

*Toxoplasma gondii* RHΔhxgprt strain was maintained by growth in confluent primary human foreskin fibroblasts (HFFs) in Dulbecco’s modified Eagle’s medium (DMEM; Invitrogen, Carlsbad, CA) with 10% fetal bovine serum (FBS; HyClone, Logan, UT), 2 mM glutamine, 100 U/ml penicillin, and 100 μg/ml streptomycin (cDMEM) at 37°C in 5% CO2. The HFFs are fully deidentified and therefore do not constitute human subjects research.

*Plasmodium falciparum* strain 3D7 was routinely cultured in de-identified human erythrocytes from the Stanford Blood Center at 3% hematocrit at 37°C, 5% CO_2_ and 1% O_2_. The culture medium (termed complete RPMI) consisted of RPMI-1640 (Sigma, St. Louis, MO) supplemented with 25 mM HEPES, 50 mg/L hypoxanthine (Sigma), 2.42 mM sodium bicarbonate, and 4.31 mg/ml Albumax II (Gibco, Waltham, MA).

### Parasite preparation

*Toxoplasma* tachyzoites were released from heavily infected monolayers of HFFs by mechanical disruption of the monolayers using disposable scrapers and passage through a 25-gauge syringe. The parasites were added to fresh monolayers of HFFs, and 18-20 hours post-infection, were washed two times with Endo buffer (EB) (45 mM potassium sulfate, 106 mM sucrose, 10 mM magnesium sulfate, 20 mM Tris buffer pH 7.2, 5 mM glucose and 0.35% bovine serum albumin). The HFFs monolayers were scraped and passage through a 27-gauge syringe and tachyzoites were released into fresh EB at room temperature. Tachyzoites were pelleted and resuspended in fresh EB.

To stimulate the parasites, tachyzoites were washed two times with cold Hank’s balanced salt solution (HBSS) without calcium, magnesium and phenol red (Corning, Corning, NY), supplemented with 1 mM MgCL_2_, 1 mM CaCl_2_, 10 mM NaHCO_3_ and 20 mM HEPES, pH 7. The HFFs monolayers were scraped and passage through a 27-gauge syringe and tachyzoites were released into fresh cold HBSS. Tachyzoites were pelleted and resuspended in fresh cold HBSS. Note that all steps were done on ice. Calcium-ionophore (A23187, Sigma) at a final concentration of 1 μM was added to the sample at room temperature followed by chemical fixation for conoid protrusion assay or plunge freezing of unfixed sample for cryogenic electron tomography.

*Plasmodium* merozoites were isolated as previously described by Boyle et al. [41], with some modifications. *P. falciparum* was synchronized to the schizont stage using sequential rounds of 5% sorbitol (Sigma) and magnet purification using LS columns (Miltenyi Biotec) and a MACS magnet. On the day of harvest, 120 ml of *P. falciparum* culture at 10-20% parasitemia was subjected to MACS magnet purification to isolate schizonts away from uninfected erythrocytes, yielding ~5×10^8^ schizonts. The schizonts were resuspended in 40 ml fresh complete RPMI with 10 μM E-64 (Sigma), a cysteine protease inhibitor that prevents merozoite release from schizonts by inhibiting rupture, but does not adversely affect merozoites [41, 42]. After a 5 hour incubation at 37°C, the cells were pelleted at 1500 rpm for 5 minutes and supernatant removed completely. The schizont pellet was resuspended in 6 ml of PBS supplemented with 2% FBS, 5 mM MgCl_2_, and 50 μg/ml DNase I, and transferred to a syringe connected to a 1.2 μm filter. To release the merozoites, pressure was applied to the syringe plunger evenly until all liquid was expelled. The flow-through was applied to an LS column on a MACS magnet to remove hemozoin, and then centrifuged at 4000 rpm for 10 minutes, followed by two washes in PBS. The merozoites were quantified using a hemocytometer, and resuspended at 5×10^7^/ml or 5×10^6^/ml.

### Conoid protrusion

Tachyzoites were prepared as described above and treated for 0.5-2 minutes with A23187 or incubated for 10 minutes in EB at room temperature. Tachyzoites were then fixed for 20 minutes with Formaldehyde and concentrated by centrifugation. The pellet was resuspended in 30 μl PBS and left to dry on a glass coverslip. Protruded and non-protruded parasites were counted using the oil 100x magnification under Phase condition. The average number of protruded parasites was determined by counting 60 parasites for each condition for three independent experiments. Data are presented as mean values ± SD from three independent experiments.

### Western blot

Tachyzoites were prepared as described above and treated for 0, 2, or 5 minutes. The tachyzoites were pelleted from the soup and both fractions were boiled for 5 min, separated by SDS-PAGE, and transferred to polyvinylidene difluoride (PVDF) membranes. Secreted TgMIC2 was detected by incubation of membrane with mouse anti-TgMIC2 antibody (6D10, ascites) and followed by incubation with horseradish-peroxidase (HRP) conjugated goat anti-mouse IgG. The levels of HRP were detected using an enhanced chemiluminescence (ECL) kit (Pierce). SAG1 levels were used to control for the concentration of parasites within the samples using rabbit anti-SAG1 followed by staining with secondary HRP-conjugated goat anti-rabbit IgG antibody and detecting the levels of HRP as described above.

### Cryogenic electron tomography and reconstruction

Glow discharge lacey carbon EM grids were mounted on a manual plunger, loaded with parasite suspension mixed with 10 nm gold fiducials (EMS), blotted from the back side using Whatman paper #5, and plunged into liquid ethane at near liquid nitrogen temperature.

The stimulated *Toxoplasma* tachyzoites and the purified *Plasmodium* merozoites were imaged using a Talos Arctica electron microscope (Thermo Fisher) equipped with a field emission gun operated at 200kV, a Volta Phase Plate [43], an energy filter (Gatan) operated at zero-loss and a K2 Summit direct electron detector (Gatan). Upon phase plate alignment and conditioning, tilt series of the parasites were recorded at 39,000x at pixel size 3.54 Å using Tomo4 software with bidirectional acquisition schemes, each from −60° to 60° with 2° or 3° increment. Target defocus was set to −1.0 μm. The K2 camera was operated in dose fractionation mode recording frames every 0.6s. Every 2 or 3 tilt series, a new spot on the phase plate was selected. The phase shift spans a range from 0.2-0.8π. The total dose was limited to 70-90 e/Å2.

The non-stimulated tachyzoites were imaged using a Titan Krios electron microscope (Thermo Fisher) equipped with a field emission gun operated at 300kV, a Volta Phase Plate, an energy filter (Gatan) operated at zero-loss and a K2 Summit direct electron detector (Gatan). Tilt series of the parasites were recorded at 26,000x at pixel size 3.46 Å. Tilt series were recorded using Tomo4 software with bidirectional acquisition schemes, each from −60° to 60° with 2° or 3° increment. The phase shift spans a range from 0.2-0.3π. Target defocus was set to −1.0 μm. The total dose was limited to 60-100 e/Å2. The movie frames were motion-corrected using motionCor2 [44], and the resulting micrographs are compiled into tilt series. For screening, tilt series alignment and reconstruction was performed automatically using the tomography pipeline in EMAN2 [45]. For analysis, tilt series alignment and reconstruction was performed using IMOD [46, 47]. The reconstructed tomograms are the result of multiple frozen grids that were imaged over several multiple-day sessions.

### Mixed-Scale Dense Networks and iterative augmentation

We trained on 20 initial annotated images from 7 different tomograms to build an initial network. In each of the 7 tomograms we picked a slice that included the most information on our features of interest, and manually annotated it with two to three adjacent slices (one or two above and one below). Each slice was fully annotated with different labels for the different features and an additional label to exclude the area outside of the cell from the training data. We then added an additional 43 annotated images to the training set by allowing the network to extract the major features, and then manually made corrections: the initial predictions were quite accurate, requiring only limited corrections. These corrections, for the most part, were in the form of correcting missing areas of AVs, rhoptries and IMTs structures and were made on 3-6 adjacent slices, picked by their relevance to the biological question. The network was then trained on the entire, annotated set.

In each case, we trained an MSD network with hyperparameters that are typically used in other studies as well: a width of one, a depth of 100, and dilations uniformly chosen between one and ten, resulting in a network with 49574 learnable parameters (compared to over 31 million learnable parameters in a standard U-Net). The network input consisted of the 2D cryo-ET slice to be annotated, with four adjacent slices (two above and two below) used as additional input. During training, the manual annotations were used as training target, minimizing the Dice loss. To artificially increase the amount of available training data, the data was augmented using random 90° rotations and flips. All NN computations were performed using the code accompanying [25], which is GPU-accelerated and available under an open-source license (https://github.com/dmpelt/msdnet). Training was terminated when no significant improvement in the loss was observed, which was typically after a few days.

### Meta data extraction from annotated tomograms

Each of the following quantifications used all tomograms that resolved the feature of interest. Top views were not included in the analysis since the relevant features are not well resolved.

Connected components, surfaces, volumes, and centers of mass were calculated using the image processing toolbox within MATLAB version R2020b. All distances between structures were calculated in three dimensions unless otherwise specified. Average vesicle curvature values shown in Fig. 5b were calculated using the central 11 slices (~±7.5 nm) of each vesicle. These measurements were then averaged for each class of vesicle to produce the plots in Fig. 5b. Vesicle volume measurements were scaled by a factor of 1.33 to account for the missing wedge. This factor was estimated from central slices of IMT-associated AVs that were assumed to be spherical. Incorporation of this factor produces values closer to the true volume of the vesicles than a sum of annotated voxels alone.

## Data availability

The tomograms of the *Toxoplasma* apical end are deposited to EMDB with the accession code D_1000266664 for the tomogram of the stimulated tachyzoite, and D_1000266665 for the tomogram of the non-stimulated tachyzoite.

## Acknowledgments

We thank all members of our respective research groups for always helpful input and especially Melanie Espiritu for growth of fibroblasts and Dr. Michael F. Schmid for particularly insightful comments during analysis of the data. We thank Christina Hueschen, Alex Dunn, Rachel Stadler, and Gary Ward for helpful discussions. We thank Chan Zuckerberg Biohub Intercampus Team Award for supporting this research. Other support includes NIH grants: S10OD021600, P41GM103832, P01GM121203, DP2HL137186, Stanford Maternal Child Health Research Institute, and the Center for Advanced Mathematics for Energy Research Application (CAMERA), jointly funded by The Office of Advanced Scientific Research (ASCR) and the Office of Basic Energy Sciences (BES) within the DOE’s Office of Science at the Lawrence Berkeley National Laboratory, under contract number DE0AC02-5CH11231. L.S-Z. was supported in part by BARD, the United States - Israel Binational Agricultural Research and Development Fund, Vaadia-BARD Postdoctoral Fellowship Award No. FI-582-2018, and Stanford School of Medicine Dean’s Postdoctoral Fellowship. P.D.D was supported in part by the National Institute of General Medical Sciences grant R35-GM118067. D.M.P. was supported in part by The Netherlands Organization for Scientific Research (NWO), project number 016.Veni.192.235.

## Supplementary material

**Figure S1. Stimulation with calcium ionophore (A23187) induces conoid protrusion and microneme secretion at room temperature. (A)** Percentage of conoid protrusion in tachyzoites stimulated with 1 μM of A23187 (in HBSS) or non-stimulated (Endo buffer). Data are represented as mean ± SD with HBSS+A23187 giving 80.1±4.6% (N=196) vs. with Endo buffer giving 7.4±2.1% (N=816). **(B)** Microneme secretion was analyzed by western blot, showing the cellular and predicted cleaved forms of MIC2. Tachyzoites were harvested into HBSS with calcium ionophore (HBSS+A23187) or non-stimulating buffer (Endo buffer). MIC2 was detected using mouse anti-MIC2 (6D10, ascites) antibody. Antibodies against *Toxoplasma* surface antigen 1 (rabbit anti-SAG1) were used to control for equal loading. Size markers (kDa) are shown on the left of the blot.

**Figure S2. Additional observations regarding the rhoptries and anterior AVs in the AC. (A)** A tomographic slice of a representative AC from an ionophore-stimulated tachyzoite showing a constriction of the rhoptry’s neck (left) and highlighted in red (right). The solid rectangles show the continuation of the rhoptry neck on a different z-slice of the dashed area. Scale bar, 200 nm. A movie of the complete tomogram of (A) showing the constriction of the rhoptry is available in S3 Movie. **(B)** A tomographic slice showing two AVs (numbered arrows) in the apex of an AC from an ionophore-stimulated tachyzoite. Scale bar, 200 nm. **(C)** As for (B) but slices from a different tomogram showing three discrete anterior AVs. **(D)** The left panel is a tomographic slice from an ionophore-stimulated tachyzoite showing an anterior AV interacting with two rhoptries. Scale bar, 200 nm. The middle (unannotated) and right (structures highlighted in colors) panels are a zoomed-in view of the square from the left panel showing the tip of the rhoptries (red), and the AV (yellow). The points of interaction between the AV and the rhoptries are marked by white arrowheads. Scale bar, 50 nm. **(E)** as for (D) except the anterior AV is interacting with just one rhoptry.

**Figure S3. NN derived annotations are similar to ones made by expert human annotators and improves with rounds of training.** Tomographic slices (from the same reconstructed tomogram) that were either used **(A)** or not used **(B)** in the training data (left panels), their manual annotation by an expert (middle panels), and the NN annotation after training #1 (right panels). The annotation is showing the AVs (yellow), rhoptries and micronemes (red), and microtubules (blue). Scale bar, 200 nm. Differences in the annotation of the AVs, rhoptry, and IMT between the NN and an expert person are pointed by the arrows. **(C)** The percentage agreement in the annotation between two expert persons (P1 vs. P2) or each of the expert persons and the NN after training #3 (P1/2 vs. NN) for the AVs, micronemes/rhoptries, and microtubules (MTs; subpellicular and IMTs). Data are represented as mean ± SD (N=3). Agreement percentage is based either on each pixel error (“pixel-wise”) or on connected components. Annotated pixels were grouped into connected components if they shared any edge or corner in two-dimensions and connected components from different annotations were considered to be mutually identified if they shared 25% or more pixels in common. In no case was there a statistically significant difference between any of the pair-wise comparisons for a given parameter.

**Figure S4. NN derived annotations improves with rounds of training.** A tomographic slice that was not used in the training data and its annotation by the NN. The NN annotation (overlaid on the tomographic slice) showing the AVs (yellow), rhoptries and micronemes (red), and microtubules (blue) after each of 3 rounds of training. Note the small but marked improvement in the annotation accuracy of the AVs, rhoptry, and IMT with each additional training as pointed by the arrows. Note that a different slice from the same tomogram was used in training #3. Scale bar, 200 nm.

**Figure S5. A schematic for the curvature calculation for the anterior and IMT-associated AVs.** The curvature around the vesicles was measured for each pixel on the perimeter of the vesicle by fitting a circle to that pixel ±5 adjacent pixels on the periphery. The absolute value of the curvature was taken to be the inverse of the circle radius. If the center of the fit circle was found to be on the interior of the AV then it was given a positive value of curvature and a negative value if on the exterior of the AV. The average curvature was then calculated for every 10 degrees in both directions (Ψ), starting from the shortest distance between each vesicle’s COM and either the rhoptry tip or the IMT, for anterior **(A)** and IMT-associated vesicles **(B)**, respectively.

**Figure S6. More characterization of the IMTs and its associated AVs. (A)** A tomographic slice showing a vesicle posterior to the IMTs (arrowhead). The solid rectangle shows the basal part of the IMT (arrow) on a different z-slice of the dashed rectangle. Scale bar, 200 nm. **(B)** The length of the shorter and longer microtubule in each IMT pair. Each point represents data from one tomogram, and the dashed line represents the X=Y relation. **(C)** The IMT’s length in stimulated (Stim.; mean ± SD =356.5±60.3 nm median=343.2 nm, N=36) and non-stimulated (Non-Stim.; mean ± SD =304.1±32.8 nm median=308.4 nm, N=6) tachyzoites. **(D)** The IMT’s curvature in stimulated (Stim.; mean ± SD =0.7±0.4 X10^-4^ nm^-1^ median=0.6X10^-4^ nm^-1^, N=36) and non-stimulated (Non-Stim.; mean ± SD =1.1±0.8 X10^-4^ nm^-1^ median=0.9X10^-4^ nm^-1^, N=6) tachyzoites. **(E)** The distance between the anterior-most AV’s COM and the tip of the closest IMT. **(F)** A histogram showing the length distribution for the densities between IMT-associated AVs. **(G)** A histogram showing the minimal distance between neighboring AVs (membrane to membrane) that had observed densities between them (“connected”) and without such densities (“unconnected”).

**Figure S7. A density is covering both the anteriorly located and IMT-associated AVs. (A)** Right panel is a tomographic slice showing a density partially surrounding the anteriorly located AV in an ionophore-stimulated tachyzoite. Scale bar, 200 nm. Middle panel is zoomed in view of the square shown in the left panel. Scale bar, 50 nm. Right panel is a repeat of the middle panel with the density surrounding the AV highlighted in magenta. **(B)** As for (A) except from a non-stimulated tachyzoite. **(C,D)** As for (A) and (B), respectively, except showing IMT-associated AVs.

**Figure S8. More characterization of the density covering both the anteriorly located and IMT-associated AVs. (A)** Tomographic slices showing two anterior AVs in the same tomogram (left panel; arrowheads) from a stimulated tachyzoite. Scale bar, 200 nm. Middle panels are the top views (average of 12 successional slices from orthogonal sectioning planes labeled in red dashed lines in the left panel) of the AVs (middle left) and the density (middle right) covering them. Right panel is as for the middle right panel except it highlights in color the rosette-like density covering the AV. Scale bar, 50 nm. **(B)** and **(C)** Tomographic slices showing an IMT-associated AV (arrowhead) from a stimulated tachyzoite. Scale bar, 200 nm. Right panel is the average of 12 (B) or 3 (C) successional slices of the density covering the IMT associated AV. Insets are zoomed-in views of the squares, showing that the densities have a rosette-like pattern. Scale bar, 50 nm. **(D)** As for (C) except from a non-stimulated tachyzoite.

**Movie S1: The 3-D organization of the apical complex of non-stimulated *Toxoplasma gondii* tachyzoite.**

**Movie S2: The 3-D organization of the apical complex of ionophore stimulated *Toxoplasma gondii* tachyzoite.**

**Movie S3: Tomogram showing the constriction of the rhoptry.**

**Movie S4: 3-D annotation of the AV of ionophore stimulated tachyzoite.**

