## Supplementary material for "Coupling cryo-electron tomography with mixed-scale dense neural networks reveals re-organization of the invasion machinery of *Toxoplasma gondii* upon ionophore-stimulation"

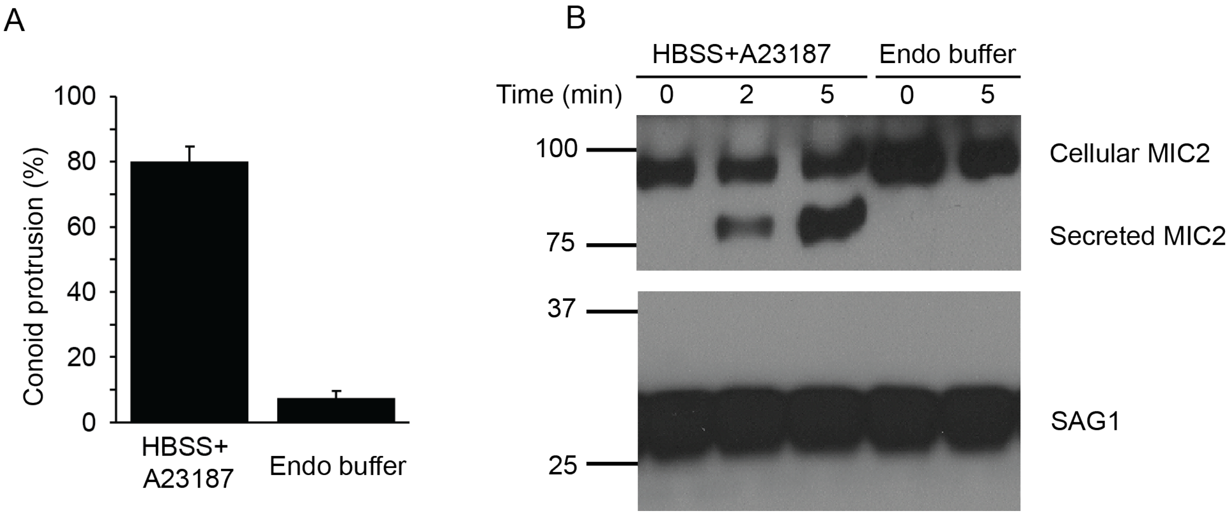


**S1 Fig. Stimulation with calcium ionophore (A23187) induces conoid protrusion and microneme secretion at room temperature.**

(**A**) Percentage of conoid protrusion in tachyzoites stimulated with 1 µM of A23187 (in HBSS) or non-stimulated (Endo buffer). Data are represented as mean ± SD with HBSS+A23187 giving 80.1±4.6% (N=196) vs. with Endo buffer giving 7.4±2.1% (N=816). (**B**) Microneme secretion was analyzed by western blot, showing the cellular and predicted cleaved forms of MIC2. Tachyzoites were harvested into HBSS with calcium ionophore (HBSS+A23187) or non-stimulating buffer (Endo buffer). MIC2 was detected using mouse anti-MIC2 (6D10, ascites) antibody. Antibodies against *Toxoplasma* surface antigen 1 (rabbit anti-SAG1) were used to control for equal loading. Size markers (kDa) are shown on the left of the blot.

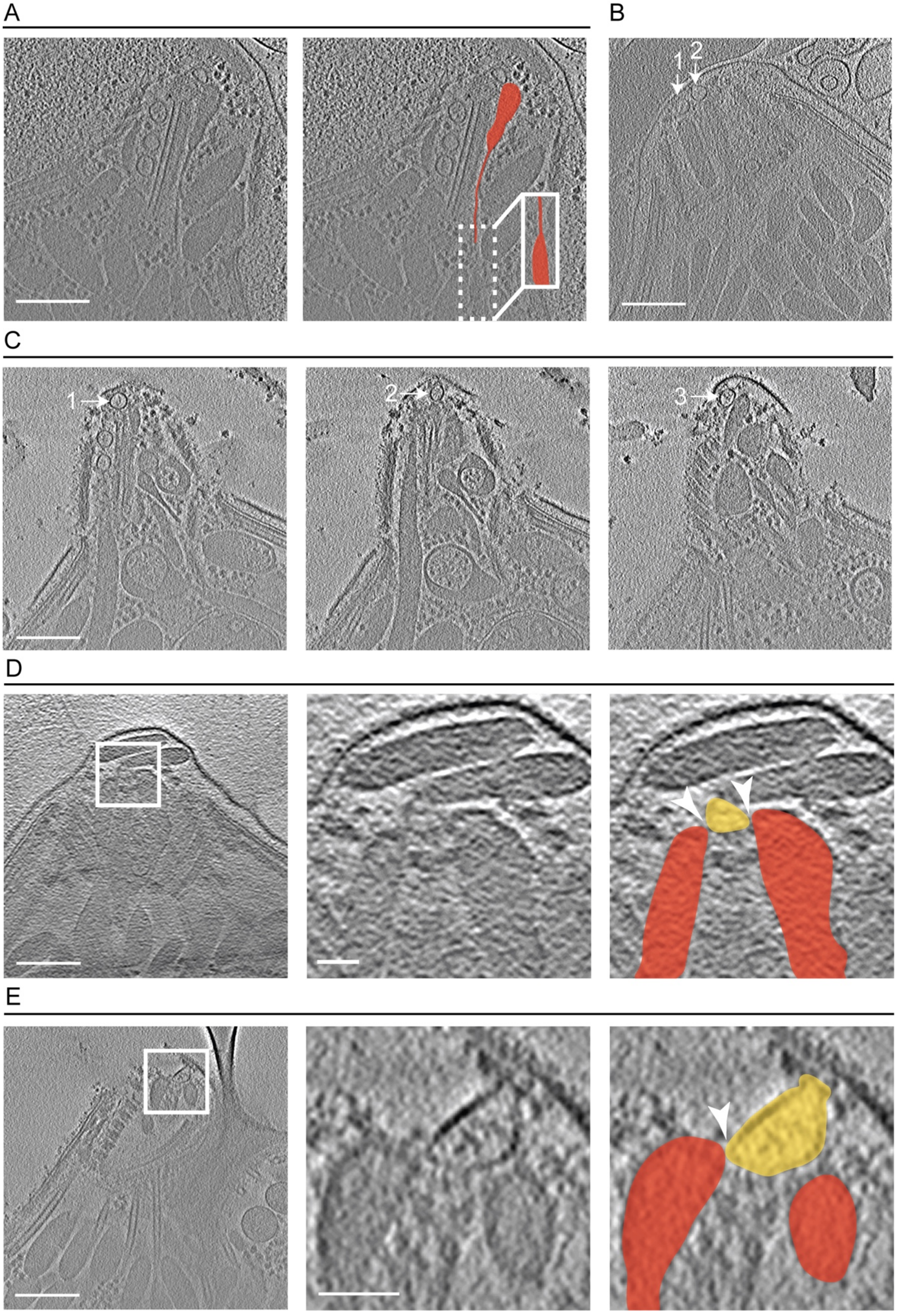
A
B
1
2
C
1
2
3
D
E

**S2 Fig. Additional observations regarding the rhoptries and anterior AVs in the AC.**
(**A**) A tomographic slice of a representative AC from an ionophore-stimulated tachyzoite showing a constriction of the rhoptry's neck (left) and highlighted in red (right). The solid rectangles show the continuation of the rhoptry neck on a different z-slice of the dashed area. Scale bar, 200 nm. A movie of the complete tomogram of (A) showing the constriction of the rhoptry is available in S3 Movie. (**B**) A tomographic slice showing two AVs (numbered arrows) in the apex of an AC from an ionophore-stimulated tachyzoite. Scale bar, 200 nm. (**C**) As for (B) but slices from a different tomogram showing three discrete anterior AVs. (**D**) The left panel is a tomographic slice from an ionophore-stimulated tachyzoite showing an anterior AV interacting with two rhoptries. Scale bar, 200 nm. The middle (unannotated) and right (structures highlighted in colors) panels are a zoomed-in view of the square from the left panel showing the tip of the rhoptries (red), and the AV (yellow). The points of interaction between the AV and the rhoptries are marked by white arrowheads. Scale bar, 50 nm. (**E**) as for (D) except the anterior AV is interacting with just one rhoptry.

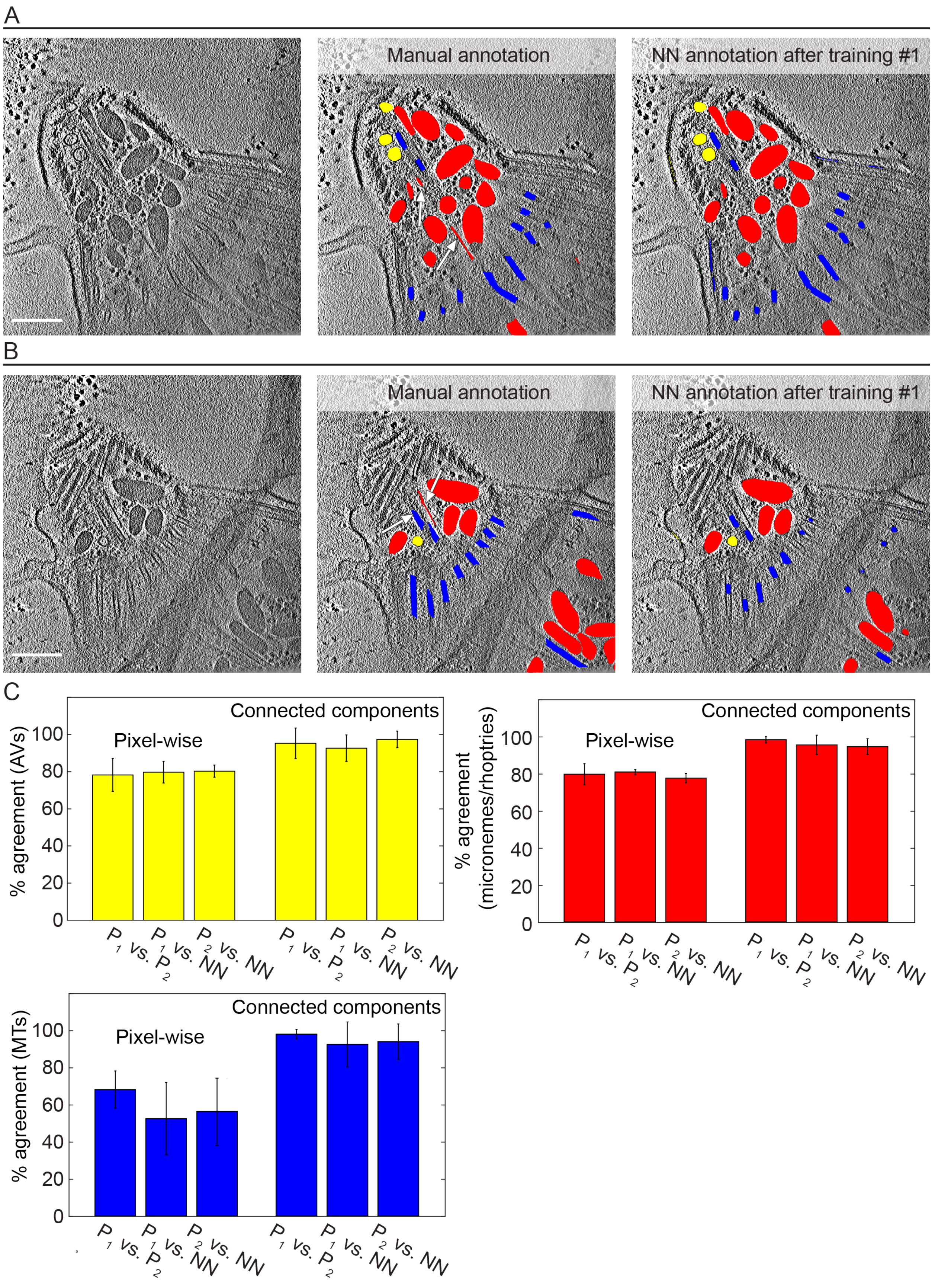
A
Manual annotation
NN annotation after training #1
B
Manual annotation
NN annotation after training #1
C
% agreement (AVs)
Pixel-wise
Connected components
% agreement (micronemes/rhoptries)
Pixel-wise
Connected components
% agreement (MTs)
Pixel-wise
Connected components
P1 vs. P2
P1 vs. NN
P2 vs. NN
0
20
40
60
80
100

**S3 Fig. NN derived annotations are similar to ones made by expert human annotators and improves with rounds of training.**

Tomographic slices (from the same reconstructed tomogram) that were either used (**A**) or not used (**B**) in the training data (left panels), their manual annotation by an expert (middle panels), and the NN annotation after training #1 (right panels). The annotation is showing the AVs (yellow), rhoptries and micronemes (red), and microtubules (blue). Scale bar, 200 nm. Differences in the annotation of the AVs, rhoptry, and IMT between the NN and an expert person are pointed by the arrows. (**C**) The percentage agreement in the annotation between two expert persons ($P_1$ vs. $P_2$) or each of the expert persons and the NN after training #3 ($P_{1/2}$ vs. NN) for the AVs, micronemes/rhoptries, and microtubules (MTs; subpellicular and IMTs). Data are represented as mean ± SD (N=3). Agreement percentage is based either on each pixel error ("pixel-wise") or on connected components. Annotated pixels were grouped into connected components if they shared any edge or corner in two-dimensions and connected components from different annotations were considered to be mutually identified if they shared 25% or more pixels in common. In no case was there a statistically significant difference between any of the pair-wise comparisons for a given parameter.

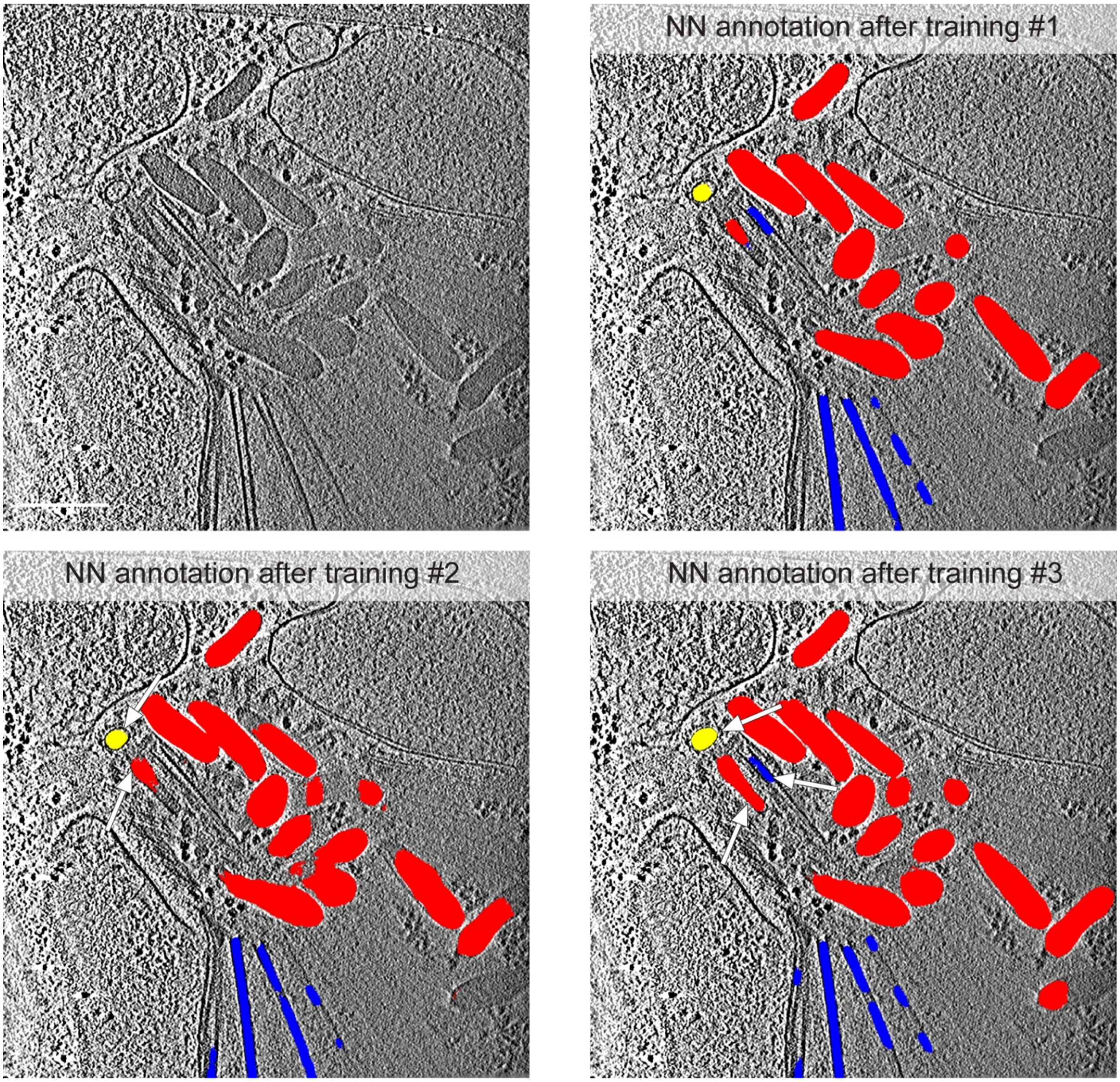


**S4 Fig. NN derived annotations improves with rounds of training.**
A tomographic slice that was not used in the training data and its annotation by the NN. The NN annotation (overlaid on the tomographic slice) showing the AVs (yellow), rhoptries and micronemes (red), and microtubules (blue) after each of 3 rounds of training. Note the small but marked improvement in the annotation accuracy of the AVs, rhoptry, and IMT with each additional training as pointed by the arrows. Note that a different slice from the same tomogram was used in training #3. Scale bar, 200 nm.

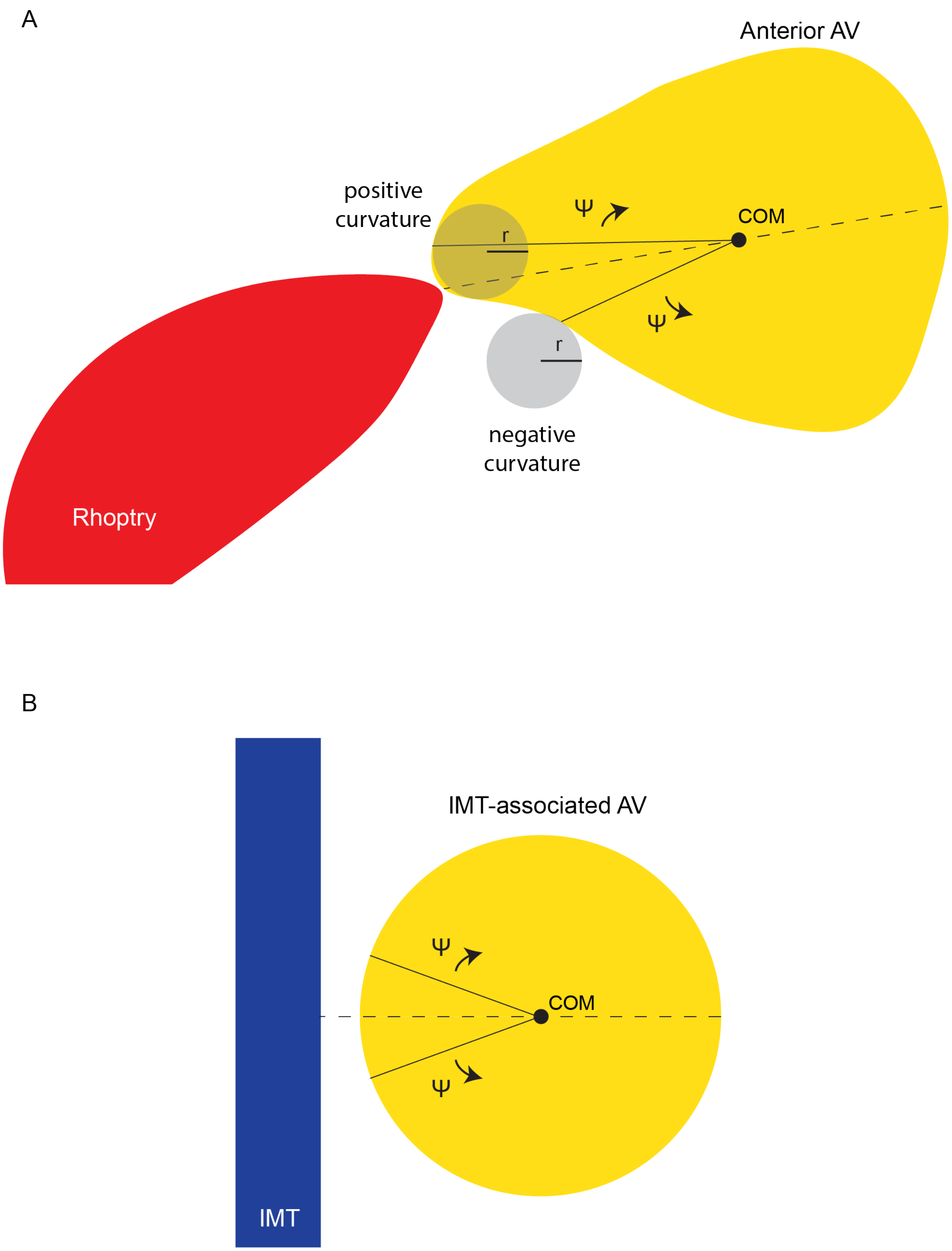
A
Anterior AV
positive
curvature
Ψ
r
COM
Ψ
r
negative
curvature
Rhoptry
B
IMT-associated AV
Ψ
COM
Ψ
IMT

**S5 Fig. A schematic for the curvature calculation for the anterior and IMT-associated AVs.** The curvature around the vesicles was measured for each pixel on the perimeter of the vesicle by fitting a circle to that pixel ±5 adjacent pixels on the periphery. The absolute value of the curvature was taken to be the inverse of the circle radius. If the center of the fit circle was found to be on the interior of the AV then it was given a positive value of curvature and a negative value if on the exterior of the AV. The average curvature was then calculated for every 10 degrees in both directions (Ψ), starting from the shortest distance between each vesicle's COM and either the rhoptry tip or the IMT, for anterior (**A**) and IMT-associated vesicles (**B**), respectively.

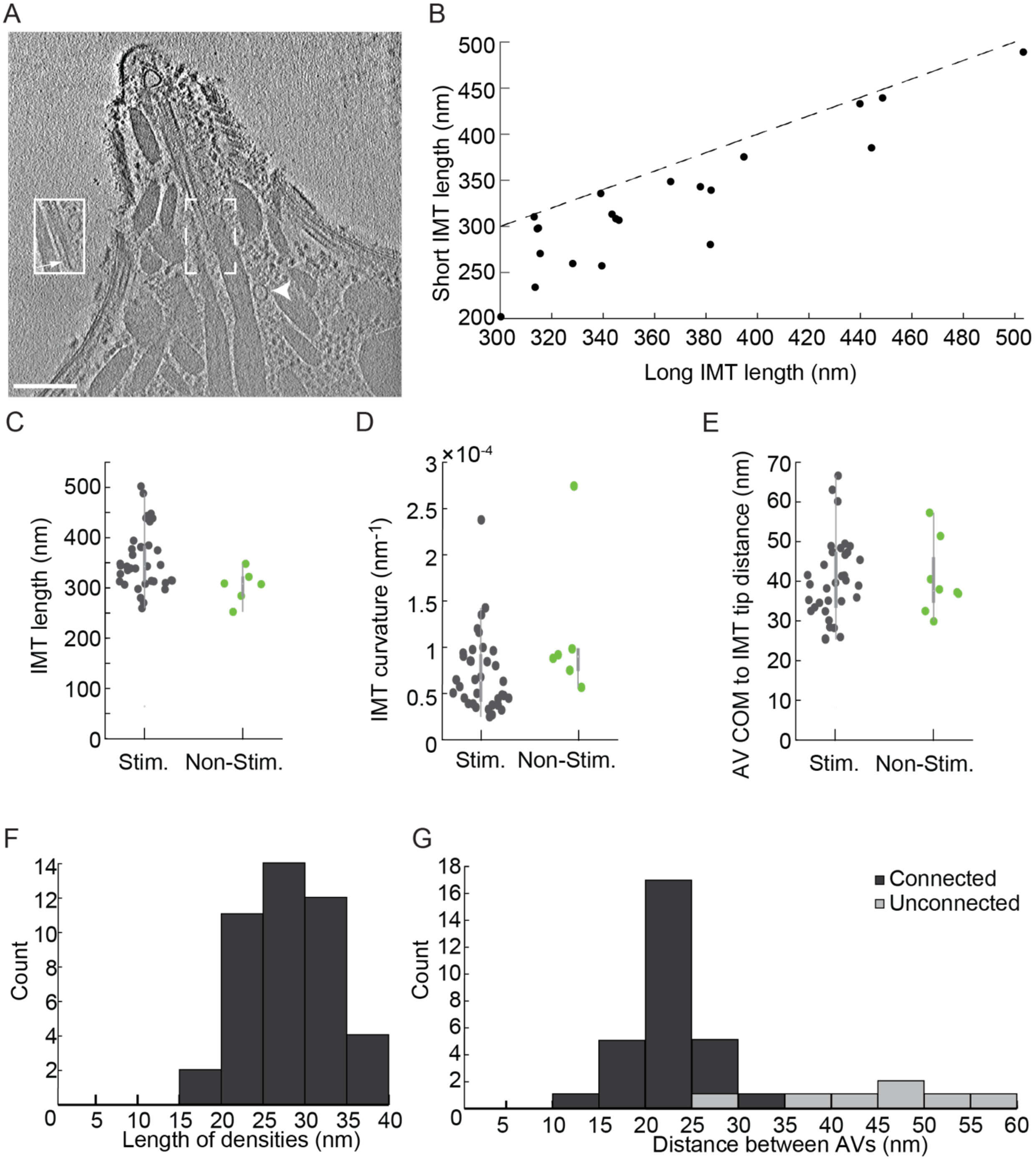


**S6 Fig. More characterization of the IMTs and its associated AVs.**

(**A**) A tomographic slice showing a vesicle posterior to the IMTs (arrowhead). The solid rectangle shows the basal part of the IMT (arrow) on a different z-slice of the dashed rectangle. Scale bar, 200 nm. (**B**) The length of the shorter and longer microtubule in each IMT pair. Each point represents data from one tomogram, and the dashed line represents the X=Y relation. (**C**) The IMT's length in stimulated (Stim.; mean ± SD =356.5±60.3 nm median=343.2 nm, N=36) and non-stimulated (Non-Stim.; mean ± SD =304.1±32.8 nm median=308.4 nm, N=6) tachyzoites. (**D**) The IMT's curvature in stimulated (Stim.; mean ± SD =0.7±0.4 X10$^{-4}$ nm$^{-1}$ median=0.6X10$^{-4}$ nm$^{-1}$, N=36) and non-stimulated (Non-Stim.; mean ± SD =1.1±0.8 $X10^{-4}$ $nm^{-1}$ median=0.9$X10^{-4}$ $nm^{-1}$, N=6) tachyzoites. (**E**) The distance between the anterior-most AV's COM and the tip of the closest IMT. (**F**) A histogram showing the length distribution for the densities between IMT-associated AVs. (**G**) A histogram showing the minimal distance between neighboring AVs (membrane to membrane) that had observed densities between them ("connected") and without such densities ("unconnected").

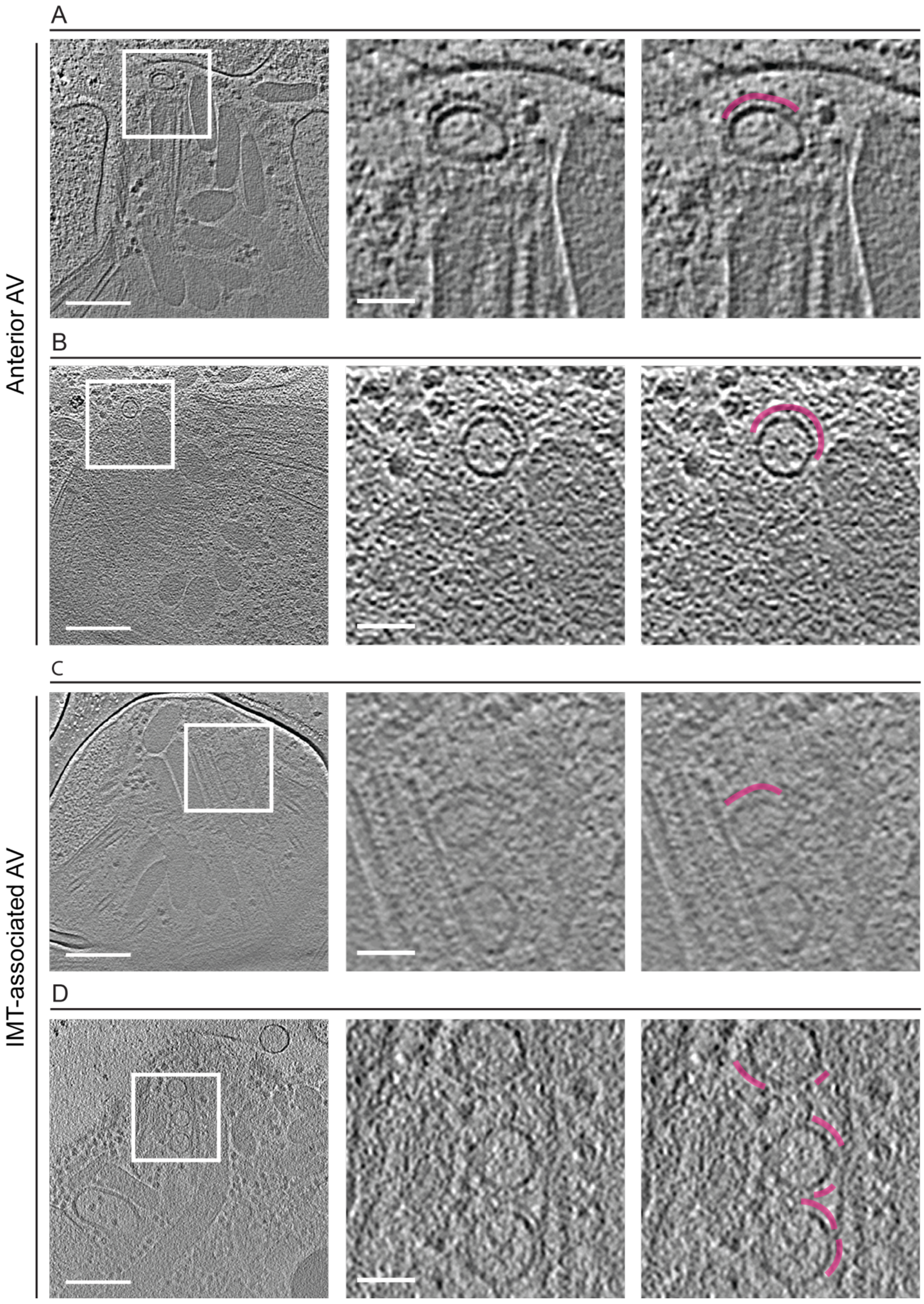
A
B
C
D
Anterior AV
IMT-associated AV

**S7 Fig. A density is covering both the anteriorly located and IMT-associated AVs.**
(**A**) Right panel is a tomographic slice showing a density partially surrounding the anteriorly located AV in a stimulated tachyzoite. Scale bar, 200 nm. Middle panel is zoomed in view of the square shown in the left panel. Scale bar, 50 nm. Right panel is a repeat of the middle panel with the density surrounding the AV highlighted in magenta. (**B**) As for (A) except from a non-stimulated tachyzoite. (**C,D**) As for (A,B, respectively) except showing IMT-associated AVs.

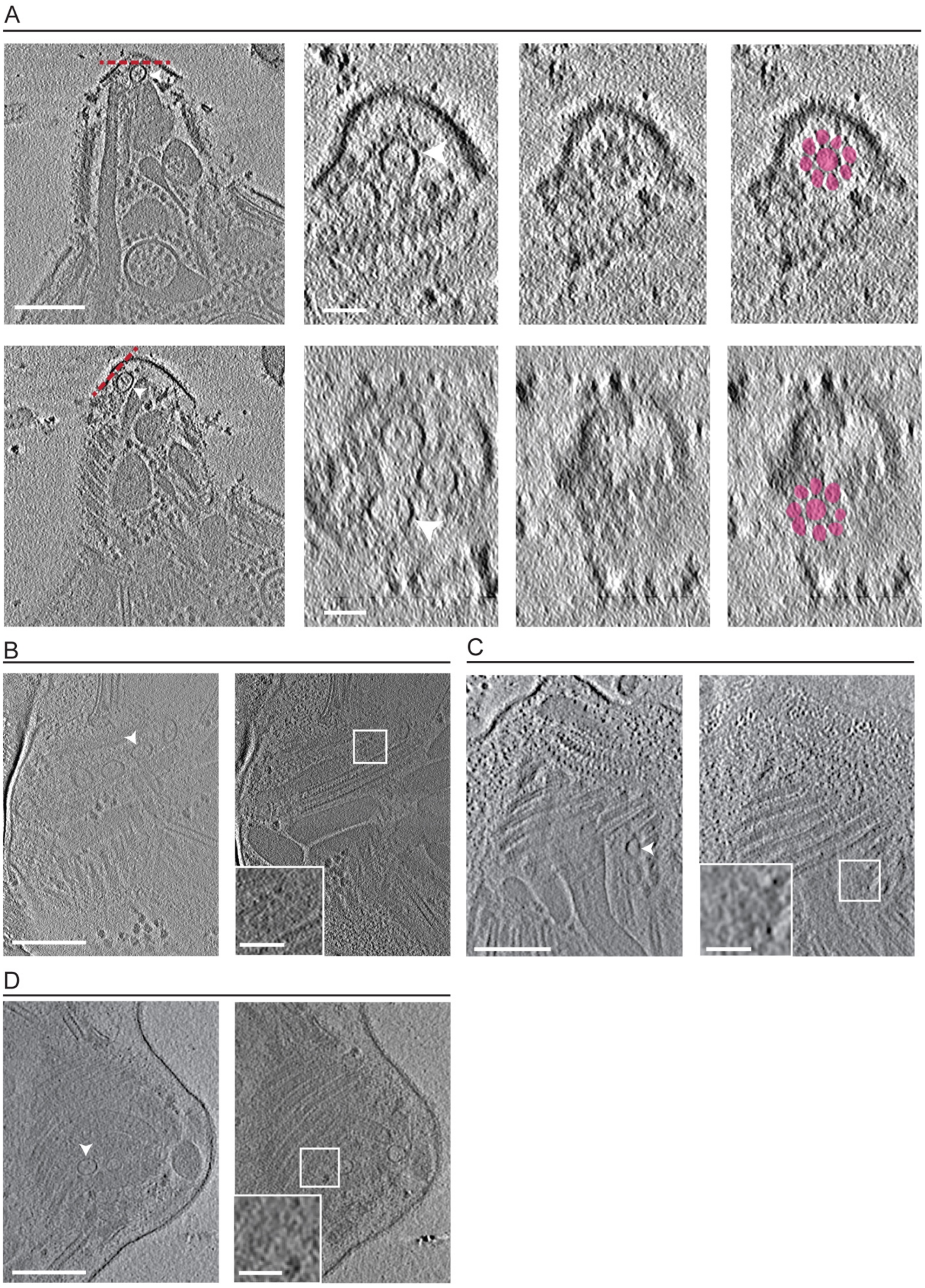
A
B
C
D

**S8 Fig. More characterization of the density covering both the anteriorly located and IMT-associated AVs.**
(**A**) Tomographic slices showing two anterior AVs in the same tomogram (left panel; arrowheads) from a stimulated tachyzoite. Scale bar, 200 nm. Middle panels are the top views (average of 12 successional slices from orthogonal sectioning planes labeled in red dashed lines in the left panel) of the AVs (middle left) and the density (middle right) covering them. Right panel is as for the middle right panel except it highlights in color the rosette-like density covering the AV. Scale bar, 50 nm. (**B**) and (**C**) Tomographic slices showing an IMT-associated AV (arrowhead) from a stimulated tachyzoite. Scale bar, 200 nm. Right panel is the average of 12 (B) or 3 (C) successional slices of the density covering the IMT associated AV. Insets are zoomed-in views of the squares, showing that the densities have a rosette-like pattern. Scale bar, 50 nm. (**D**) As for (C) except from a non-stimulated tachyzoite.

S1 Movie. The 3-D organization of the apical complex of non-stimulated *Toxoplasma gondii* tachyzoite.

S2 Movie. The 3-D organization of the apical complex of ionophore stimulated *Toxoplasma gondii* tachyzoite.

S3 Movie. Tomogram showing the constriction of the rhoptry.

S4 Movie. 3-D annotation of the AV of ionophore stimulated tachyzoite.
